# Optimizing copy number profiles for tumor cell phylogeny inference from low-coverage single-cell sequencing data

**DOI:** 10.64898/2026.10.05.756871

**Authors:** Naeem Ahmed, Samson Weiner, Carter Payne, Mukul S. Bansal

## Abstract

**Motivation:** Accurate reconstruction of tumor cell phylogenies is crucial for understanding cancer evolutionary dynamics, inferring intra-tumor heterogeneity, and designing effective and personalized cancer treatment strategies. One of the most promising and cost-effective approaches for reconstructing tumor phylogenies for large numbers of cells is to use copy number alteration (CNA) data obtained from *low-coverage single-cell DNA sequencing* of sampled tumor cells. However, error and uncertainty in inferred copy number profiles (CNPs), resulting from low sequence coverage, can have a drastic negative effect on the reconstruction accuracy of tumor cell phylogenies.

**Results:** In this work, we show that careful selection of *bin size*, a critical tuning parameter for inferring CNPs, is necessary to maximize the accuracy of tumor cell phylogeny inference. Specifically, we (i) use three previously published low-coverage single-cell DNA sequencing datasets from real tumors to demonstrate that even small changes in bin size can have a drastic impact on downstream tumor phylogeny reconstruction, (ii) show that this effect persists even when using simulated low-coverage datasets generated under idealized conditions with minimal noise, and (iii) propose a simple computational approach to select bin sizes that maximize tumor phylogeny accuracy. Our experiments span a wide range of bin sizes and account for the impact of various factors including coverage-level, phylogeny reconstruction method, and CNP inference method, and explicitly account for phylogenetic reconstruction uncertainty. Our evaluation of the proposed computational approach for bin size selection shows that it can significantly improve tumor cell phylogeny reconstruction accuracy compared to baseline methods.

**Availability:** Freely available from https://github.com/Tanveer14/BinSelector.

## 1 Introduction

To understand cancer progression and intra-tumor heterogeneity (ITH) at a fine-grained level, it is necessary to detect mutations in individual tumor cells and infer a phylogenetic tree that describes the evolutionary relationships between individual cells and subclones [2, 11, 19, 21]. Advancements in single-cell DNA sequencing (scDNA-seq) have been pivotal in enabling both mutation detection and tumor phylogeny inference at the single-cell level [10,21]. Because of the characteristic low coverage of tumor scDNA-seq data (typically 0.02x to 0.1x), CNAs, which occur frequently and are often large enough to be detectable from such data, have become a widely used phylogenetic marker for tumor cell phylogeny inference [1, 3, 8, 20, 21, 26, 27, 30, 33, 35, 37]. In a typical phylogenetic analysis, copy number profiles (CNPs) are first inferred for the individual sequenced cells and then a tumor cell phylogeny is constructed using the CNPs. Both these tasks are computational in nature, and the reliability of the reconstructed tumor cell phylogeny depends on the resolution and accuracy of the computed CNPs. Indeed, error in the CNPs has a strong negative effect on the accuracy of reconstructed tumor cell phylogenies [35].

Given the importance of reliable CNP inference from raw scDNA-seq data, both for tumor cell phylogeny reconstruction and many other applications, considerable effort has been made to improve the methodologies underpinning this task [9, 16, 17, 25, 31, 32]. Since raw scDNA-seq data are notoriously noisy and sparse, single-cell CNP inference methods typically count reads within large fixed-size genomic bins (typically 100 kbp - 5 Mbp [6, 17]) to reduce the effects of variable amplification and sequence sampling [7]. Most CNP inference methods, including Ginkgo [7], SCOPE [31], CHISEL [36], SeCNV [25], and SCICoNE [9], are based on this idea and use fixed-size bins. The specific choice of *bin size* has important implications for CNA detection and CNP inference, and the task of selecting an appropriate bin size is typically left to the user. Selecting larger bin sizes can reduce variance in the read count signals, potentially making detection of large-scale CNAs more reliable, but provides only a coarse-grained, low-resolution view of CNAs. On the other hand, selecting smaller bin sizes makes it possible to detect smaller CNAs but also makes CNP inference more susceptible to noise and biases in scDNA-seq. Thus, for tumor cell phylogeny reconstruction, larger bin sizes yield CNPs that provide less, but potentially higher-quality, information for phylogeny construction. In contrast, smaller bin sizes yield CNPs that provide more, but lower-quality, information for phylogeny construction. In general, for any particular experiment, the optimal choice of bin size for phylogeny construction may depend on various factors including sequence coverage, degree of sequencing bias, distribution of CNA event sizes, etc.

In this work, we address two closely related questions: First, how sensitive are reconstructed tumor cell phylogenies to changes in bin size? And second, given scDNA-seq data from tumor cells, how can we select a bin size that maximizes the accuracy of tumor cell phylogeny inference from that data? These questions are motivated by our previous analyses of two published tumor scDNA-seq datasets, T10 and T16 [21], where we observed that even modest changes in bin size can significantly affect reconstructed tumor cell phylogenies and lead to altered inferences regarding the number and composition of clonal populations within a tumor [35]. For example, when we analyzed the 100-cell T16 cancer dataset (consisting of 52 cells from a primary breast tumor and 48 cells from its associated liver metastasis from patient T16), changing the bin size from 100 kbp to 250 kbp resulted in the inference of different clone compositions. The resulting CNP heatmaps are shown in Supplementary Figure S1. In particular, the 250 kbp bins support the presence of an additional clonal population, whereas the 100 kbp bins group those cells with an existing clonal population. Such inconsistencies, if ignored, can be problematic and have important medical implications.

### Related work

Although the importance of this parameter has been acknowledged in literature and the question of ‘what should the bin size be?’ has been asked [17, 23, 29], few solutions to this problem have been presented. For example, [23] offers that bin size intrinsically depends on the interest and type of analysis, but provides no framework for arriving at a value. Tu et al. [29] used downscaled sequencing data from single cells to assess the sensitivity of CNP inference to sequencing depth and bin size and suggested that 0.75× sequencing depth with a 250 kbp bin size should work well in most cases. Some approaches that optimize bin size, use variable bin sizes, or are bin-free have also been developed [21, 32, 38]. One of the earliest approaches for CNP inference used variable-sized bins [21], where each bin contained a roughly equal number of reads. However, that approach still requires an input parameter for specifying the average reads-per-bin and has a number of other drawbacks addressed by later methods. HiScanner [38] takes a unique approach of leveraging the presence of allelic dropout (ADO), a technical artifact of scDNA-seq where one allele fails to be amplified. Dropout results in read count fluctuations that closely resemble a copy number deletion, but these effects are dampened with sufficient bin size. The idea behind HiScanner is to learn the distribution of ADO (amplicon) sizes, which is specific to the amplification protocol, using a hidden markov model (HMM) learned on normal cells. Then, the bin size is set to be the 95th percentile of dropout sizes. However, HiScanner needs bulk sequencing data, in addition to scDNA-seq data, to estimate ADO. SCNV [32] adapts a bin-free segmentation approach designed for bulk sequencing [28] to single-cells. That original approach [28] uses generalized likelihood ratio statistics to detect points in the genome where the read intensity shifts in the target sample, as compared to a normal cell reference. The number of change points is chosen using point-wise approximate Bayesian confidence intervals. SCNV adapts this approach to single-cells [32].

Overall, despite the above methods, and even approaches that use trees to help improve CNA inference [9, 18], there has been no systematic attempt (to the best of our knowledge) to assess the impact of bin size on tumor cell phylogeny reconstruction or to optimize bin size with the specific goal of yielding CNPs that maximize tumor cell phylogeny accuracy.

### Our contributions

We make three key contributions: First, we use three real low-coverage single-cell sequencing datasets to demonstrate that even small changes in bin size can have a drastic impact on downstream tumor phylogeny reconstruction. These findings hold across a wide range of bin sizes and across different coverage levels, phylogeny reconstruction methods, and CNP inference methods. Importantly, we find that when two very different tumor cell phylogeny inference methods (MEDICC2 [8] and DICE-star [35] in our experiments) are applied to the same bin size, the resulting phylogenies are often much more similar to each other than when the same method is applied to different (even neighboring) bin sizes. This implies that the drastic differences in tumor cell phylogenies for different bin sizes are not a result of tree reconstruction error, uncertainty, or lack of phylogenetic signal in the data. Second, we use an extensive simulation study to show that this effect persists even for simulated low-coverage datasets generated under idealized conditions free from possible confounding factors. The persistence of this effect on simulated datasets well-suited for tumor cell phylogeny reconstruction further supports our previous findings on real datasets and reinforces the importance of careful bin size selection. Third, we propose a simple computational method to select bin sizes that maximize tumor phylogeny accuracy. We evaluate our method on the simulated datasets, which have known ground-truth tumor cell phylogenies, and find that it is more effective than other baseline approaches for bin size selection. Overall, this work spotlights the importance of bin size to tumor phylogeny reconstruction from CNPs inferred using low-coverage scDNAseq data, and demonstrates the feasibility of easily optimizing bin size to maximize phylogenetic accuracy even in the absence of any additional information or ground truth.

## 2 Materials and Methods

In this section, we describe, in order, the real and simulated datasets used in this work, the computational pipeline used for inferring CNPs, our methodology for assessing the impact of bin size changes on reconstructed phylogenies, the proposed method for bin size selection, and our approach for evaluating the method.

### 2.1 Description of datasets

#### Real datasets

We used three previously published low-coverage scDNA-seq datasets: CRC2, T3, and T5. The CRC2 dataset consists of 42 cells sampled from a colorectal cancer patient [12]. The mean coverage of CRC2 is 0.033x (SRA accession number: SRP074289). The T3 and T5 datasets consist of 152 and 128 tumor cells, respectively, sampled from triple-negative breast cancer (TNBC) patients [6]. Their mean coverages are 0.046x and 0.041x, respectively (SRA accession number: SRP064210). All coverages were calculated after alignment and MAPQ [15] thresholding.

#### Simulated datasets

We used CNAsim [34] to generate ground-truth tumor cell phylogenies and simulated sequencing reads from sampled tumor cells. These reads were then used for inferring CNPs, which, in turn, were subsequently used for reconstructing tumor cell phylogenies. To focus specifically on the effect of bin size changes on reconstructed tumor cell phylogenies, we chose simulation settings that minimized the effect of possible confounding factors on CNP inference accuracy. In particular, our simulation settings did not allow dropout events or amplification biases common in single-cell sequencing data. The sequencing error rate was also kept at a relatively low 0.02. The most relevant simulation parameter for CNP inference is sequence coverage. Based on the coverages of real scDNA-seq datasets, we simulated datasets with coverages 0.02x and 0.03x. All simulated datasets consisted of 100 sampled tumor cells, of which 10% are specified to be normal cells, and 10 replicates were simulated for each of the two coverage levels. Thus, the simulated datasets consist of 20 different scDNA-seq datasets. Most other simulation parameters were left at their default values. Details on specific commands and parameter settings used appear in the Supplement.

### 2.2 CNP Inference

We applied the same processing and CNA inference pipeline to the simulated and real datasets. For each cell, we used the tool BWA (Burrows-Wheeler Aligner) [13] to align the paired-end reads to the human reference genome hg38. We then used Samtools [14] to process the aligned reads and obtain the BAM files used by the CNP inference methods. We used two different CNP inference methods, SCOPE [31] and SeCNV [25]. SCOPE is a state-of-the-art method that leverages shared breakpoint signals across cells for improved segmentation and a normalization model with direct ploidy estimation. SeCNV is a more recent method that uses a local Gaussian kernel to capture pairwise similarities between bins along the genome and then partitions the genome into segments by minimizing structural entropy. Both methods take as input a user-specified bin size.

For this study, we generated CNPs using the following 11 bin sizes: 200, 400, 600, 800, 1000, 1500, 2000, 3000, 4000, 5000, and 6000 kbp. On some real or simulated datasets, certain bin sizes resulted in one or both CNP inference methods being unable to compute CNPs. When this occurred for real datasets, we attempted to use adjacent bin sizes (e.g., 300 kbp instead of 200 kbp). When this occurred for simulated datasets, we either used only the successful replicates (out of 10 total per setting) for that bin size or, if all replicates failed, we skipped that bin size entirely. For both SCOPE and SeCNV, normal cells were explicitly specified.

### 2.3 Assessing impact of bin size changes on phylogenetic reconstruction

We used the computed CNPs to infer phylogenetic trees using DICE-star [35] and MEDICC2 [8], two state-of-the-art methods for tumor cell phylogeny reconstruction. The primary goal of this phylogenetic analysis is to better understand how sensitive reconstructed tumor cell phylogenies are to changes in bin size. The most straightforward approach for assessing the impact of bin size on reconstructed tumor cell phylogenies would be to calculate phylogenetic distances between phylogenies reconstructed from CNPs of different bin sizes. However, phylogenetic inference is often intrinsically uncertain, reflecting methodological biases (e.g., different objective functions), insufficient phylogenetic signal in the input, or limitations of the phylogenetic search procedures (typically based on local search heuristics) used by the phylogeny construction methods. Thus, variation in reconstructed tumor cell phylogenies across different bin sizes must be interpreted with caution since not all phylogenetic differences can be attributed to a change in bin size.

The standard approach to account for phylogenetic reconstruction uncertainty is to compute multiple candidate reconstructions based on the same input, either by applying the same method multiple times on the same input or, more commonly, by using techniques such as bootstrapping [4] involving statistical resampling of the input. However, we found these standard approaches to be inapplicable to the current problem. In particular, both DICE-star and MEDICC2 produced the same phylogeny when run multiple times on the same input CNP, and bootstrapping is inapplicable for CNPs since copy numbers for adjacent bins are not independent and the ordering of copy numbers is important. To circumvent the above issue, we devised the following approach:

1. Apply two different tumor cell phylogeny reconstruction methods to each CNP for each bin size. We use the methods DICE-star and MEDICC2 since they are highly dissimilar to each other and utilize different objective functions.
2. Calculate phylogenetic distances between corresponding DICE-star and MEDICC2 phylogenies for each input CNP (same bin size). We refer to these distances as *intra-bin* distances. We use normalized Robinson-Foulds distance (NRFD), defined below, to calculate phylogenetic distances.
3. Calculate phylogenetic distances (NRFD) between all pairs of DICE-star (or MEDICC2) phylogenies reconstructed from CNPs of the 11 different bin sizes. We refer to these distances as *inter-bin* distances. Observe that, unlike in the previous step, the same phylogeny reconstruction method (either DICE-star or MEDICC2) is used for computing these inter-bin distances. This ensures that inter-bin distances are not unnecessarily inflated due to methodological differences.
4. Compare the resulting inter-bin and intra-bin distances to assess if phylogenetic differences across different bin sizes can indeed be attributed to changes in bin size.

To calculate phylogenetic distances between tumor cell phylogenies, we use the standard (unrooted) *normalized Robinson-Foulds distance (NRFD)* [24]. NRFD takes values between 0 and 1 and represents the fraction of non-trivial splits (leaf-set bipartitions) that are present in only one of the two phylogenetic trees being compared. Thus, lower NRFD values indicate that the topologies of the two trees being compared are more similar to each other, with an NRFD of 0 implying identical tree topologies.

The above approach makes it possible to distinguish between the effect of phylogenetic reconstruction uncertainty (lack of sufficient phylogenetic signal) from the effect of changing bin size since, without sufficient phylogenetic signal in the input CNPs, DICE-star and MEDICC2 would be unable to converge upon similar intra-bin tumor cell phylogenies given their highly disparate objective functions. For example, if the intra-bin distance (NRFD between the DICE-star and MEDICC2 phylogenies) is approximately 0.4 for a 200 kbp bin size CNP as well as for a 400 kbp bin size CNP, but the inter-bin distance (NRFD between DICE-star phylogenies) between a 200 kbp bin size CNP and a 400 kbp bin size CNP is 0.6, then this suggests that changing the bin size from 200 kbp to 400 kbp results in a greater change in the inferred phylogeny than can be explained by phylogenetic reconstruction uncertainty alone.

We applied this approach to the different simulated and real datasets described previously. By default, we used SCOPE [31] to infer CNPs and DICE-star phylogenies to calculate inter-bin distances. For completeness and additional validation, we also explored the effect of using a different CNP inference method, SeCNV [25], as well as the effect of using the MEDICC2 phylogenies for inter-bin distances.

For simulated datasets, all reported inter-bin and intra-bin are averaged over the 10 replicate datasets for each sequence coverage level. We note that, in some cases, the reported values are averaged over less than 10 replicates. This generally occurred when the CNP inference method used was unable to construct CNPs for all 10 replicates for that bin size. In a small number of cases, intra-bin NRFDs could only be computed for some of the replicates due to failed MEDICC2 runs. In our experiments, MEDICC2 was executed with at least 120 GB of RAM available, but this was insufficient for some of the MEDICC2 runs. Finally, in certain rare cases, some inter-bin NRFDs could not be computed at all since there was no overlap between the sets of replicates for which CNPs could be successfully inferred for the two bin sizes involved in that inter-bin NRFD calculation.

### 2.4 A simple computational approach for bin size selection

We now describe our proposed computational method for selecting the bin size. The method involves the following five steps:

1. Compute CNPs for a wide range of bin sizes. We recommend using the set of bin sizes used/tested in this work (200, 400, 600, 800, 1000, 1500, 2000, 3000, 4000, 5000, and 6000 kbp).
2. Compute a tumor cell phylogeny for each bin size. In this work, we use DICE-star by default and also consider MEDICC2, but a different tumor cell phylogeny construction method could be used instead.
3. Compute all pairwise phylogenetic distances between the tumor cell phylogenies. In this work, we use NRFD for phylogenetic distances.
4. For each bin size *b*, compute the average distance from the phylogeny of *b* to the phylogenies of all other bin sizes. We denote this average pairwise distance by *APD*(*b*).
5. Select the bin size *b* that minimizes *APD*(*b*).

Put simply, the method selects the bin that corresponds to the median phylogeny. The method is based on the following simple insight: Assuming the input CNP data are phylogenetically informative, the tumor cell phylogenies constructed from the CNPs of different bin sizes each represent noisy/error-prone variants of the true phylogeny. Thus, as long as sufficiently many, well-spaced bin sizes are used, the median phylogeny should represent a robust estimate of the central tendency of the data. As we show later, this simple approach works well with our chosen set of eleven bin sizes. We note that these eleven bin sizes were chosen independently, prior to evaluating this computational approach. For convenience, we refer to this method as the *bin-selector* method in the remainder of this manuscript.

### 2.5 Evaluating performance of bin-selector and other bin size selection methods

We evaluated the performance of the bin-selector method and alternative approaches using the previously described simulated datasets. These simulated datasets have known ground truth tumor cell phylogenies, making it possible, for each individual dataset, to rank the different bin sizes by accuracy. We use this ground-truth ranking to evaluate bin size selection methods. We also used NRFD between the estimated and ground-truth tumor cell phylogenies to compute regret scores.

We define the two metrics used below.

#### Normalized Rank

For each execution of the method, we calculate the *normalized rank* of the bin size chosen by the bin size selection method to be the rank of the chosen bin in the ground-truth ranking divided by the total number of bin sizes for which phylogenies could be computed for that specific scDNA-seq dataset. In most cases, the normalizing factor is 11, since we use 11 different bin sizes. However, SCOPE and SeCNV sometimes failed to produce CNPs for some bin sizes for some datasets, and MEDICC2 sometimes failed to compute tumor cell phylogenies within the specified resource limits, resulting in smaller normalization factors for such input instances.

#### Regret score

Since multiple bin sizes can sometimes lead to similar tumor cell phylogeny accuracies, we also use a regret score to measure the effectiveness of bin size selection. The regret score captures the gap between the selected outcome and the best possible outcome. We compute a normalized regret score as follows. Let *opt, chosen*, and *worst* denote the NRFD between the ground-truth tumor cell phylogeny and the phylogeny for the best bin size, phylogeny for the bin size chosen by the bin size selection method, and phylogeny for the worst bin size, respectively. Then, the (normalized) *regret score* is defined to be 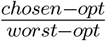.

Observe that both the above metrics take values between 0 and 1, with smaller values indicating better performance.

## 3 Results: Impact of Bin Size on Tumor Phylogenetics

We first present experimental results for the real datasets and then for the simulated datasets. For completeness and additional validation, we considered two different CNP inference methods and two different tumor cell phylogeny inference methods. These options are summarized below, with default methods highlighted in bold font.

- *CNP inference method*: **SCOPE** and SeCNV
- *Primary phylogeny reconstruction method*: **DICE-star** and MEDICC2. Here, primary method refers to the method used to compute inter-bin distances. Note that both methods are used for computing intra-bin distances.

### 3.1 Results for real datasets

Here, we demonstrate the phylogenetic impact of different bin sizes using three real low-coverage scDNA-seq datasets: CRC2, T3, and T5.

#### 3.1.1 CRC2 dataset

Table 1 (top) shows all inter-bin phylogenetic distances (NRFDs) between phylogenies computed for different bin sizes for the CRC2 dataset when using our default CNP inference method (SCOPE) and default tumor cell phylogeny inference method (DICE-star). As the table shows, phylogenies for different bin sizes are very different from one another, with NRFDs close to 1. These NRFDs remain high even for phylogenies of adjacent bin sizes; for example, the phylogenies for bin sizes 400 and 600 kbp show an NRFD of 0.795. We also find that NRFDs between phylogenies of adjacent bin sizes are generally slightly smaller than those for very different bin sizes. These results demonstrate the sensitivity of tumor cell phylogenies to bin size. Importantly, these observations remain unchanged when the alternative inference methods, SeCNV for CNP inference and MEDICC2 for tumor cell phylogeny inference, are used instead. These additional results are shown in Supplementary Tables S1 and S2. Next, we investigated if this observed sensitivity of tumor cell phylogenies to bin size is a result of phylogenetic inference error/uncertainty. Figure 1(a) shows the corresponding intra-bin phylogenetic distances (NRFD between the DICE-star and MEDICC2 phylogenies for the same bin size) as well as inter-bin NRFDs between phylogenies for adjacent bin sizes. As the figure shows, for all bin sizes, the intra-bin NRFDs are markedly smaller than the NRFDs between phylogenies for adjacent bin sizes. For example, the intra-bin NRFD for 600 kbp bin size is 0.564 but the average inter-bin NRFD for the phylogeny of bin size 600 kbp with the phylogenies of its two adjacent bin sizes (400 and 800 kbp) is much higher at 0.795. This is an important finding and implies that the drastic differences in tumor cell phylogenies for different bin sizes, even adjacent bin sizes, are not a result of phylogenetic reconstruction error/uncertainty or lack of phylogenetic signal in the data. Crucially, these findings again remain well-supported when the alternative inference methods SeCNV (Supplementary Figure S2) and MEDICC2 (Supplementary Figure S3) are used instead.

**Table 1.** Inter-bin distance matrices for tumor cell phylogenies inferred from CNPs of different bin sizes for the CRC2 and T3 datasets using default inference methods. Results for CRC2 appear in the top table and for T3 in the bottom table. The distance matrices are symmetric, phylogenetic distances are reported in NRFD, and all bin sizes are in kbp. CNPs were inferred using SCOPE and phylogenies were reconstructed using DICE-star. NRFDs between adjacent bin sizes appear shaded in blue. For CRC2, a bin size of 300 kbp was used instead of 200 kbp since MEDICC2 failed on the 200 kbp bin CNP, and results are not shown for bin size 5000 kbp since SCOPE failed to compute CNPs for this dataset for 5000 kbp.

**CRC2 dataset**
| Bin size | 300 | 400 | 600 | 800 | 1000 | 1500 | 2000 | 3000 | 4000 | 6000 |
| --- | --- | --- | --- | --- | --- | --- | --- | --- | --- | --- |
| 300 | 0.000 | 0.872 | 0.821 | 0.846 | 0.923 | 0.923 | 0.923 | 0.923 | 0.923 | 0.897 |
| 400 | 0.872 | 0.000 | 0.795 | 0.795 | 0.769 | 0.821 | 0.846 | 0.872 | 0.821 | 0.897 |
| 600 | 0.821 | 0.795 | 0.000 | 0.795 | 0.821 | 0.744 | 0.897 | 0.872 | 0.897 | 0.974 |
| 800 | 0.846 | 0.795 | 0.795 | 0.000 | 0.795 | 0.692 | 0.923 | 0.897 | 0.872 | 0.923 |
| 1000 | 0.923 | 0.769 | 0.821 | 0.795 | 0.000 | 0.744 | 0.872 | 0.872 | 0.872 | 0.897 |
| 1500 | 0.923 | 0.821 | 0.744 | 0.692 | 0.744 | 0.000 | 0.872 | 0.846 | 0.795 | 0.897 |
| 2000 | 0.923 | 0.846 | 0.897 | 0.923 | 0.872 | 0.872 | 0.000 | 0.846 | 0.744 | 0.923 |
| 3000 | 0.923 | 0.872 | 0.872 | 0.897 | 0.872 | 0.846 | 0.846 | 0.000 | 0.872 | 1.000 |
| 4000 | 0.923 | 0.821 | 0.897 | 0.872 | 0.872 | 0.795 | 0.744 | 0.872 | 0.000 | 0.821 |
| 6000 | 0.897 | 0.897 | 0.974 | 0.923 | 0.897 | 0.897 | 0.923 | 1.000 | 0.821 | 0.000 |

**T3 dataset**
| Bin size | 200 | 400 | 600 | 800 | 1000 | 1500 | 2000 | 3000 | 4000 | 5000 | 6000 |
| --- | --- | --- | --- | --- | --- | --- | --- | --- | --- | --- | --- |
| 200 | 0.000 | 0.884 | 0.905 | 0.946 | 0.912 | 0.925 | 0.939 | 0.966 | 0.946 | 0.952 | 0.946 |
| 400 | 0.884 | 0.000 | 0.884 | 0.932 | 0.898 | 0.912 | 0.952 | 0.946 | 0.939 | 0.946 | 0.952 |
| 600 | 0.905 | 0.884 | 0.000 | 0.884 | 0.884 | 0.918 | 0.925 | 0.912 | 0.925 | 0.939 | 0.939 |
| 800 | 0.946 | 0.932 | 0.884 | 0.000 | 0.878 | 0.905 | 0.925 | 0.891 | 0.905 | 0.939 | 0.939 |
| 1000 | 0.912 | 0.898 | 0.884 | 0.878 | 0.000 | 0.898 | 0.905 | 0.939 | 0.932 | 0.932 | 0.959 |
| 1500 | 0.925 | 0.912 | 0.918 | 0.905 | 0.898 | 0.000 | 0.925 | 0.932 | 0.898 | 0.939 | 0.952 |
| 2000 | 0.939 | 0.952 | 0.925 | 0.925 | 0.905 | 0.925 | 0.000 | 0.912 | 0.932 | 0.918 | 0.925 |
| 3000 | 0.966 | 0.946 | 0.912 | 0.891 | 0.939 | 0.932 | 0.912 | 0.000 | 0.884 | 0.857 | 0.925 |
| 4000 | 0.946 | 0.939 | 0.925 | 0.905 | 0.932 | 0.898 | 0.932 | 0.884 | 0.000 | 0.905 | 0.946 |
| 5000 | 0.952 | 0.946 | 0.939 | 0.939 | 0.932 | 0.939 | 0.918 | 0.857 | 0.905 | 0.000 | 0.925 |
| 6000 | 0.946 | 0.952 | 0.939 | 0.939 | 0.959 | 0.952 | 0.925 | 0.925 | 0.946 | 0.925 | 0.000 |

**Figure 1.**
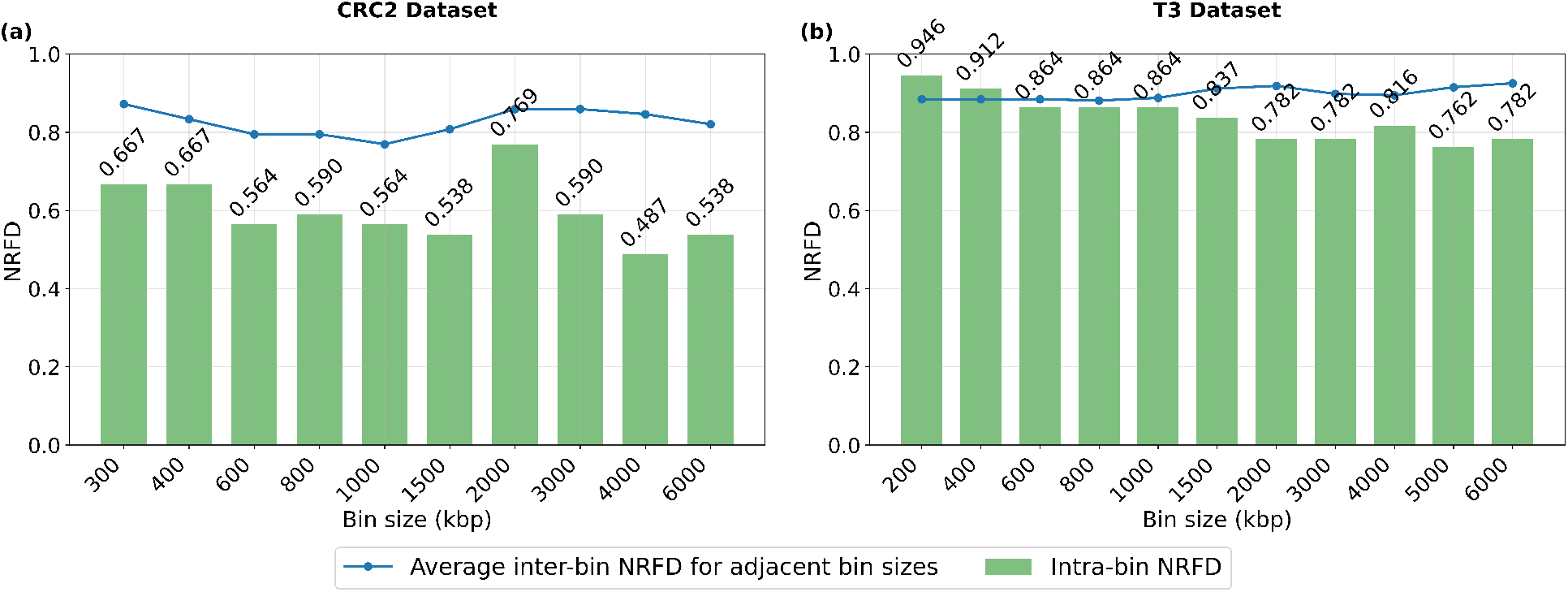
Intra-bin phylogenetic distances and inter-bin distances for adjacent bins for the CRC2 and T3 datasets. Part **(a)** shows results for CRC2 and Part **(b)** for T3. CNPs were inferred using SCOPE. For each bin size, the intra-bin distance (green bars) is the NRFD between the DICE-star phylogeny and the MEDICC2 phylogeny computed from that CNP. The inter-bin distance (blue line) reported for each bin size is the average NRFD between the DICE-star phylogeny for the current bin size and the two DICE-star phylogenies for the two adjacent bin sizes; only one adjacent bin is used for the smallest and largest bin sizes.

#### 3.1.2 T3 and T5 datasets

Analysis of the T3 and T5 datasets further support the above findings. Table 1 (bottom) and Figure 1(b) show corresponding results for the T3 dataset. We find that both intra-bin and inter-bin NRFDs are, on average, higher for the T3 dataset than for the CRC2 dataset, suggesting that T3 is a more challenging dataset for tumor cell phylogeny inference. For example, the inter-bin NRFD between the phylogenies for bin sizes 400 and 600 kbp is 0.884 and the intra-bin NRFD for 600 kbp bin size is only slightly smaller at 0.864. Still, as Figure 1(b) shows, the intra-bin NRFDs are smaller than the inter-bin NRFDs for adjacent bins across nearly all bin sizes. Specifically, intra-bin NRFDs are strictly smaller than the adjacent inter-bin NRFDs for 9 of the 11 bin sizes. Interestingly, intra-bin NRFDs are consistently lower for larger bin sizes than for smaller bin sizes, which suggests that larger bin sizes may be more appropriate for this dataset. As before, these findings are supported even when the alternative inference methods SeCNV for CNP inference (Supplementary Table S3 and Supplementary Figure S4) and MEDICC2 for tumor cell phylogeny reconstruction (Supplementary Table S4 and Supplementary Figure S5) are used; intra-bin NRFDs are strictly smaller than the adjacent inter-bin NRFDs for half of all bin sizes when SeCNV CNPs are used and for all bin sizes when MEDICC2 phylogenies are used.

The same trend is observed on the T5 dataset. T5 again shows higher intra-bin and inter-bin NRFDs than the CRC2 dataset, but, as before, the intra-bin NRFDs are strictly smaller than the inter-bin NRFDs for adjacent bins for a majority of the bin sizes. Detailed results for default inference method (Supplementary Table S5 and Supplementary Figure S6) as well as alternative inference methods (Supplementary Tables S6, S7 and Supplementary Figures S7, S8) appear in the supplement. Interestingly, on this dataset, the inter-bin NRFD for the phylogenies of 5000 and 6000 kbp bin is unexpectedly low when SCOPE CNPs are used. Specifically, this NRFD is 0.288 for DICE-star phylogenies (Supplementary Table S5) and 0.36 for MEDICC2 phylogenies (Supplementary Table S7). In contrast, the inter-bin NRFD for the phylogenies of 4000 and 5000 kbp bins is 0.936 for DICE-star phylogenies. This anomaly skews the inter-bin NRFD for adjacent bin sizes for 5000 and 6000 kbp (Supplementary Figure S6). Notably, the anomaly vanishes when SeCNV CNPs are used (Supplementary Figure S7), resulting in a clearer separation between intra-bin and inter-bin NRFDs.

### 3.2 Results for simulated datasets

Our results on the three real datasets above demonstrate that (i) reconstructed tumor cell phylogenies can be highly sensitive to changes in bin size, and (ii) even a modest change in bin size, such as using an adjacent bin size, often results in a larger change in the reconstructed tumor cell phylogeny than can be explained by phylogenetic reconstruction error or uncertainty alone. We used additional simulated datasets to assess if these findings hold even for simulated datasets simulated under idealized conditions free from possible confounding factors that could negatively impact CNP inference and tumor cell phylogeny reconstruction.

#### 3.2.1 Results for default inference methods

Results for simulated datasets using default inference methods are shown in Table 2 and Figures 2(a) and 2(b). The top table of Table 2 shows the impact of changing bin size on the resulting tumor cell phylogeny for the 0.03x sequence coverage datasets, and the bottom table for the 0.02x coverage datasets. Two important trends are immediately apparent: First, inter-bin NRFDs are much smaller for these simulated datasets than for the real datasets. This is not surprising given that simulation parameters were chosen to minimize noise and favor CNP inference and phylogenetic reconstruction, and since the CNA evolutionary models used cannot fully capture the complexity of real CNA events in tumor evolution. Second, phylogenetic differences are generally much smaller between similar bin sizes than between more disparate bin sizes. For example, for the 0.03x datasets, the average NRFD between phylogenies for bin sizes 800 and 1000 kbp is 0.281 while the NRFD for bin sizes 800 and 1500 kbp is 0.417. This, again, stands in contrast with what was observed for the real datasets, where NRFDs for adjacent bin sizes were only slightly lower than those for more disparate bin sizes. These differences point to the fact that the simulated datasets represent idealized conditions for CNP inference.

**Table 2.** Inter-bin distance matrix for tumor cell phylogenies inferred from CNPs of different bin sizes for the simulated datasets using default inference methods. Results for the 0.03x sequence coverage datasets appear in the top table and for the 0.02x coverage datasets in the bottom table. The distance matrices are symmetric, phylogenetic distances are reported in NRFD, and all bin sizes are in kbp. CNPs were inferred using SCOPE and phylogenies were reconstructed using DICE-star. NRFDs between adjacent bin sizes appear shaded in blue. All NRFDs are averaged over 10 simulated datasets.

**Simulated datasets with 0.03x sequence coverage**
| Bin size | 200 | 400 | 600 | 800 | 1000 | 1500 | 2000 | 3000 | 4000 | 5000 | 6000 |
| --- | --- | --- | --- | --- | --- | --- | --- | --- | --- | --- | --- |
| 200 | 0.000 | 0.293 | 0.304 | 0.325 | 0.371 | 0.448 | 0.528 | 0.694 | 0.786 | 0.800 | 0.800 |
| 400 | 0.293 | 0.000 | 0.293 | 0.322 | 0.361 | 0.459 | 0.530 | 0.683 | 0.790 | 0.818 | 0.820 |
| 600 | 0.304 | 0.293 | 0.000 | 0.273 | 0.323 | 0.419 | 0.497 | 0.695 | 0.789 | 0.818 | 0.809 |
| 800 | 0.325 | 0.322 | 0.273 | 0.000 | 0.281 | 0.417 | 0.482 | 0.683 | 0.781 | 0.808 | 0.796 |
| 1000 | 0.371 | 0.361 | 0.323 | 0.281 | 0.000 | 0.403 | 0.494 | 0.672 | 0.773 | 0.814 | 0.804 |
| 1500 | 0.448 | 0.459 | 0.419 | 0.417 | 0.403 | 0.000 | 0.479 | 0.672 | 0.775 | 0.805 | 0.788 |
| 2000 | 0.528 | 0.530 | 0.497 | 0.482 | 0.494 | 0.479 | 0.000 | 0.644 | 0.769 | 0.801 | 0.786 |
| 3000 | 0.694 | 0.683 | 0.695 | 0.683 | 0.672 | 0.672 | 0.644 | 0.000 | 0.766 | 0.797 | 0.838 |
| 4000 | 0.786 | 0.790 | 0.789 | 0.781 | 0.773 | 0.775 | 0.769 | 0.766 | 0.000 | 0.775 | 0.799 |
| 5000 | 0.800 | 0.818 | 0.818 | 0.808 | 0.814 | 0.805 | 0.801 | 0.797 | 0.775 | 0.000 | 0.750 |
| 6000 | 0.800 | 0.820 | 0.809 | 0.796 | 0.804 | 0.788 | 0.786 | 0.838 | 0.799 | 0.750 | 0.000 |

**Simulated datasets with 0.02x sequence coverage**
| Bin size | 200 | 400 | 600 | 800 | 1000 | 1500 | 2000 | 3000 | 4000 | 5000 | 6000 |
| --- | --- | --- | --- | --- | --- | --- | --- | --- | --- | --- | --- |
| 200 | 0.000 | 0.332 | 0.356 | 0.367 | 0.400 | 0.436 | 0.525 | 0.607 | 0.707 | 0.771 | 0.769 |
| 400 | 0.332 | 0.000 | 0.324 | 0.344 | 0.387 | 0.436 | 0.543 | 0.665 | 0.732 | 0.790 | 0.803 |
| 600 | 0.356 | 0.324 | 0.000 | 0.329 | 0.359 | 0.446 | 0.544 | 0.633 | 0.735 | 0.789 | 0.819 |
| 800 | 0.367 | 0.344 | 0.329 | 0.000 | 0.361 | 0.442 | 0.532 | 0.645 | 0.732 | 0.787 | 0.807 |
| 1000 | 0.400 | 0.387 | 0.359 | 0.361 | 0.000 | 0.436 | 0.519 | 0.650 | 0.732 | 0.795 | 0.806 |
| 1500 | 0.436 | 0.436 | 0.446 | 0.442 | 0.436 | 0.000 | 0.467 | 0.631 | 0.672 | 0.774 | 0.792 |
| 2000 | 0.525 | 0.543 | 0.544 | 0.532 | 0.519 | 0.467 | 0.000 | 0.595 | 0.682 | 0.753 | 0.792 |
| 3000 | 0.607 | 0.665 | 0.633 | 0.645 | 0.650 | 0.631 | 0.595 | 0.000 | 0.647 | 0.701 | 0.784 |
| 4000 | 0.707 | 0.732 | 0.735 | 0.732 | 0.732 | 0.672 | 0.682 | 0.647 | 0.000 | 0.713 | 0.799 |
| 5000 | 0.771 | 0.790 | 0.789 | 0.787 | 0.795 | 0.774 | 0.753 | 0.701 | 0.713 | 0.000 | 0.729 |
| 6000 | 0.769 | 0.803 | 0.819 | 0.807 | 0.806 | 0.792 | 0.792 | 0.784 | 0.799 | 0.729 | 0.000 |

**Figure 2.**
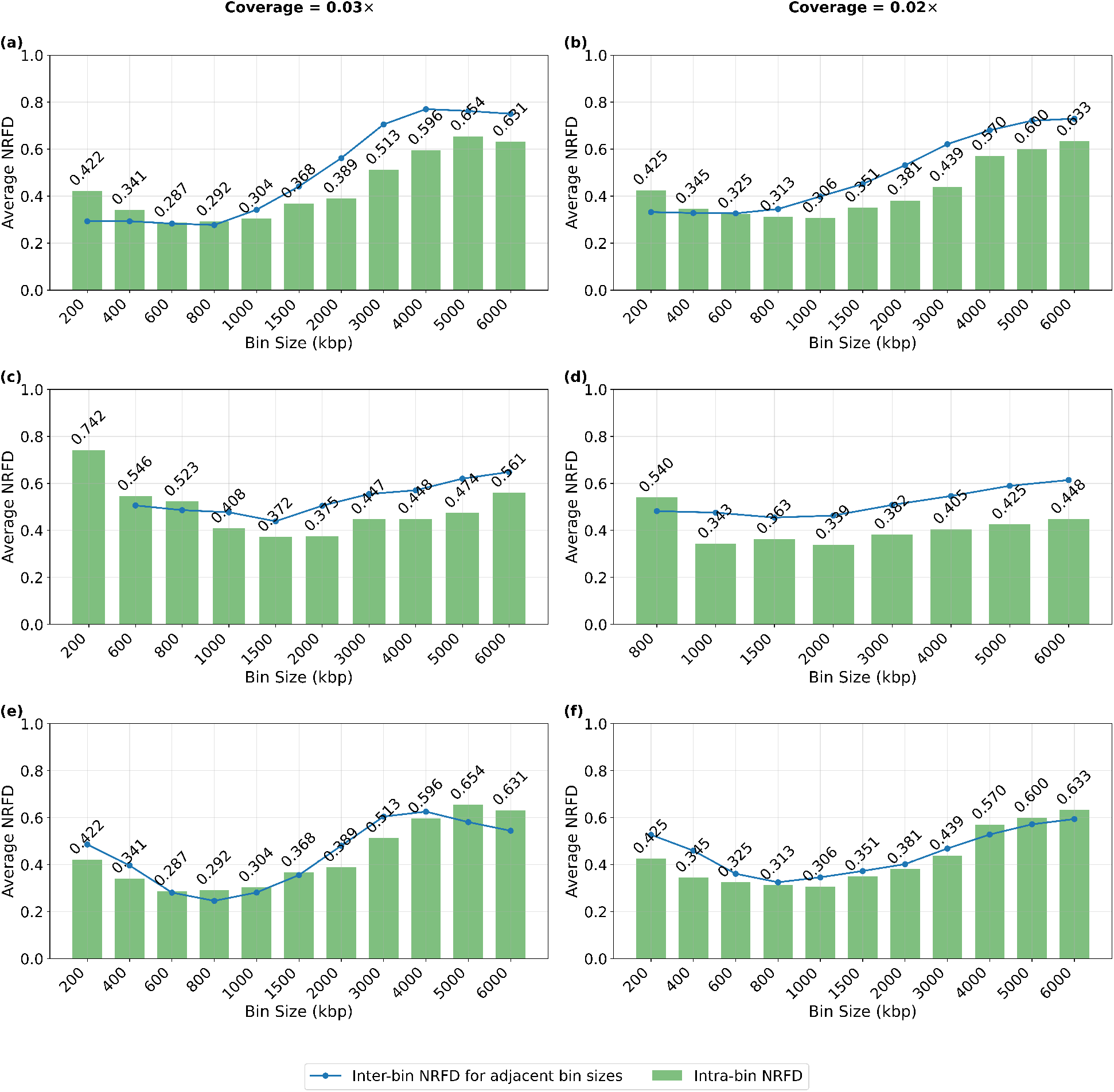
Intra-bin phylogenetic distances and inter-bin distances for adjacent bins for simulated datasets. Plots for sequence coverage 0.03x are shown in the left column and for 0.02x coverage in the right column. Plots for different CNP inference methods and different primary phylogeny reconstruction methods as organized as follows: **(a, b)** SCOPE CNPs, DICE-star phylogenies; **(c, d)** SeCNV CNPs, DICE-star phylogenies; **(e, f)** SCOPE CNPs, MEDICC2 phylogenies. For each bin size, the intra-bin distance (green bars) is the average NRFD between the DICE-star phylogeny and the MEDICC2 phylogeny computed from that CNP. The inter-bin distance (blue line) reported for each bin size is the average NRFD between the DICE-star phylogeny for the current bin size and the two DICE-star phylogenies for the two adjacent bin sizes; only one adjacent bin is used for the smallest and largest bin sizes. All NRFDs are averaged over 10 simulated datasets. Observe that several bin sizes are missing from the SeCNV plots (parts c, d). This is either because SeCNV failed to compute any CNPs for those bin sizes or because intra-bin NRFDs could not be computed for those values due to failed MEDICC2 runs.

Despite the idealized characteristics of the simulated datasets, we find that NRFDs between phylogenies reconstructed from CNPS of more disparate bin sizes remain very high. For example, the average NRFD between phylogenies constructed for bin sizes 1000 and 5000 kbp is 0.814 for the 0.03x datasets. Moreover, as Figure 2(a) for the 0.03x datasets shows, intra-bin NRFDs are smaller than inter-bin NRFDs even for adjacent bin sizes in most cases. This is consistent with previous observations made for the real datasets and clearly demonstrates the extraordinary sensitivity of phylogeny inference to small changes in bin size, even under idealized conditions free from confounding factors.

##### Effect of changing sequence coverage

The effect of changing bin size is similarly strong for the 0.02x sequence coverage datasets. As Table 2 (bottom) shows, inter-bin distances are generally a bit higher across smaller bin sizes and generally a bit lower across larger bin sizes for the 0.02x coverage datasets as compared to the 0.03x coverage datasets. Importantly, as Figure 2(b) for 0.02x coverage shows, the gap between intra-bin NRFDs and inter-bin NRFDs for adjacent bin sizes increases further, with 9 out of 11 bin sizes showing smaller intra-bin NRFDs. These results suggest that careful selection of bin size is even more important for datasets with lower sequence coverage.

##### Impact of bin size optimization

We also used the simulated datasets to directly assess the importance of optimizing bin sizes, i.e., the impact of selecting a good bin size over a bad bin size on the accuracy of the reconstructed tumor cell phylogeny. Specifically, we leveraged the availability of ground-truth phylogenies for these datasets to calculate, for each replicate, the accuracies (NRFD against the ground-truth phylogeny) of the tumor cell phylogenies reconstructed from the CNPs of different bin sizes. We then calculated the difference between the NRFDs for the most accurate reconstructed phylogeny (corresponding to the best bin size selection) and the least accurate reconstructed phylogeny (corresponding to the worst bin size selection). We refer to this difference in NRFDs as the *accuracy gap*.

For the 0.03x sequence coverage datasets the mean and median accuracy gaps are 0.525 and 0.548, respectively, and for the 0.02x coverage datasets these numbers are 0.485 and 0.494, respectively. Thus, both the 0.03x and 0.02x coverage datasets show very large accuracy gaps. This implies that selecting a good bin size can have a profound impact on the accuracy of the reconstructed tumor cell phylogeny.

#### 3.2.2 Results for alternative inference methods

##### Using SeCNV for CNP inference

We see the same overall trend for intra-bin versus inter-bin NRFDs when the CNP inference method SeCNV is used instead of SCOPE. As Figure 2(c) for the 0.03x coverage datasets shows, intra-bin NRFDs remain smaller than inter-bin NRFDs for adjacent bins for most bin sizes. As before, this separation is even more pronounced for the 0.02x coverage datasets (Figure 2(d)). Interestingly, inter-bin NRFDs remain substantially more stable across different bin size pairs overall when SeCNV CNPs are used (Supplementary Table S8) than when SCOPE CNPs are used (Table 2), more closely matching what was observed on the real datasets.

##### Using MEDICC2 for tumor cell phylogeny reconstruction

As before, these findings remain unchanged when MEDICC2 is used as the primary tumor cell phylogeny reconstruction method (Figures 2(e) and 2(f) and Supplementary Table S9) and we again see that most intra-bin NRFDs remain smaller than inter-bin NRFDs even for adjacent bins. Again, the separation between intra-bin NRFDs and inter-bin NRFDs for adjacent bins is more pronounced for the 0.02x coverage datasets (Figure 2(f)). A notable difference between results using MEDICC2 and results using our default method DICE-star is that, with MEDICC2, intra-bin NRFDs are smaller than inter-bin NRFDs for adjacent bins for the smaller bin sizes but not for the larger ones (Figures 2(e) and 2(f)), while we see the opposite trend with DICE-star (Figures 2(a) and 2(b)).

The consistency of the above findings across different sequence coverages, CNP inference methods, and tumor cell phylogeny reconstruction methods is striking and our results on simulated datasets strongly support our findings on the real datasets. Overall, these results clearly demonstrate the importance of careful bin size selection.

## 4 Results: Bin Size Selection

In this section, we evaluate the performance of the previously described bin-selector method. There are two existing methods that also consider the problem of bin size selection, HiScanner [38] and SCNV [32]. HiScanner leverages allelic dropout sizes to select bin size, with a goal of mitigating the effect of ADO as a technical artifact. However, HiScanner needs bulk sequencing data, in addition to scDNA-seq data, to estimate allelic dropout and is therefore not applicable when using scDNA-seq data alone. SCNV uses a promising, bin-free approach to produce copy number profiles. However, the software implementation of SCNV is not well documented and we were unable to get it to work despite substantial effort. We were also unable to get in touch with the developers.

Given the inapplicability of HiScanner and the difficulty with applying SCNV, we compare the performance of bin-selector to two different approaches inspired by how actual users might select a bin size for their analysis. We describe these two approaches below:

### Selecting a bin size at random

In this approach, a bin size is selected at random from among the available options. In our experiments, a bin size is chosen randomly from among the 11 available bin sizes for most instances. Fewer than 11 bin sizes were available in some instances due to failed CNP inference or MEDICC2 phylogeny inference runs. We refer to this as the *random-selection* approach.

### Selecting frequently used fixed bin sizes

Several existing studies have used bin sizes close to 500 kbp [22], [7], [31], [5]. Accordingly, we evaluated two fixed-bin-size schemes: One in which the bin size is fixed at **400 kbp** and another in which it is fixed at **600 kbp**. We refer to these two approaches as *fixed-bin-size-400* and *fixed-bin-size-600*.

In the following, we show the results of applying the bin-selector to the 0.03x and 0.02x sequence coverage simulated datasets and compare its performance against those of the random-selection, fixed-bin-size-400, and fixed-bin-size-600 approaches. For each of the 10 replicates at each coverage level, we applied all four combinations of inference methods (SCOPE+DICE-star, SCOPE+MEDICC2, SeCNV+DICE-star, SeCNV+MEDICC2), for a total of 40 input instances for each sequence coverage level.

### 4.1 Performance of the bin-selector method

Figure 3 shows the results of applying bin-selector to the 40 input instances for 0.03x sequence coverage (plots on the left) and the 40 input instances for the 0.02x coverage (plots on the right). In terms of normalized rank, we find that bin-selector almost always selects a bin size that is among the top 50% of best bin sizes (top plots of Figure 3). Specifically, it does so for 95% of the input instances for the 0.03x coverage datasets and for 90% of the input instances for the 0.02x datasets. The method identifies the best possible bin size in 40% (16 out of 40) of the 0.03x coverage input instances and in 30% (12 out of 40) of the 0.02x coverage input instances.

**Figure 3.**
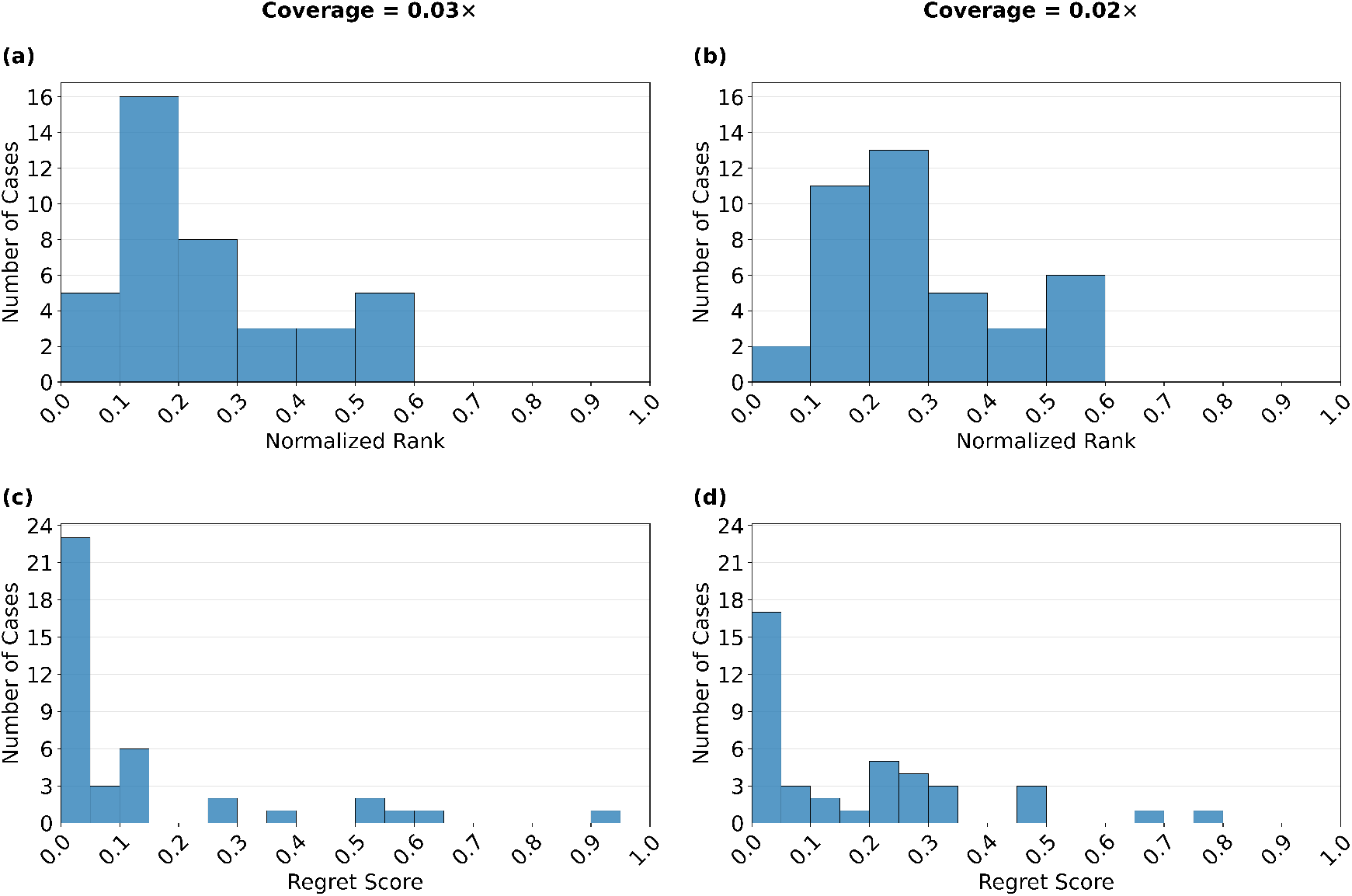
Performance evaluation of the bin-selector method. The four plots show the results of applying bin-selector to the 40 input instances for 0.03x sequence coverage simulated datasets (plots a, c) and the 40 input instances for the 0.02x coverage simulated datasets (plots b, d). The top two plots (a, b) show distributions of normalized ranks of the bins selected by bin-selector in the ground-truth ranking of bins based on the accuracies of reconstructed tumor cell phylogenies. The bottom two plots (c, d) show the distributions of (normalized) regret scores for the bins selected by bin-selector. The step size of regret scores on the x-axis is 0.05. Smaller regret scores imply better performance. Barring extreme values, x-axis intervals in all plots are left closed and right open.

Since multiple bin sizes may lead to almost equally good phylogenetic reconstruction accuracy, the regret score is a much better measure of method performance than normalized rank. The two bottom plots of Figure 3 show the distribution of regret scores for bin-selector. As those two plots show, most of the bin sizes selected by bin selector have negligible regret. For the 0.03x coverage datasets, the mean and median regret scores are only 0.133 and 0.033, respectively. These values remain low even for the more challenging 0.02x coverage datasets at 0.173 and 0.103, respectively. These results show that bin-selector is effective at selecting bin sizes that are well suited for tumor cell phylogeny reconstruction. The results also show that, as expected, the performance of the method degrades as sequence coverage decreases.

### 4.2 Comparison against alternative approaches

We compared the performance of bin-selector against the random-selection, fixed-bin-size-400, and fixed-bin-size-600 approaches. Figure 4 shows the results of this analysis. As expected, bin-selector shows far better performance than the random-selection approach (plots in top row of Figure 4). For example, for the 0.03x sequence coverage input instances, bin-selector selects a strictly better bin size than random-selection in 72.5% of cases, an equally good bin size in 20% of cases, and a worse bin size in 7.5% of cases. These values are similar for the 0.02x coverage input instances at 70%, 12.5%, and 17.5%, respectively.

**Figure 4.**
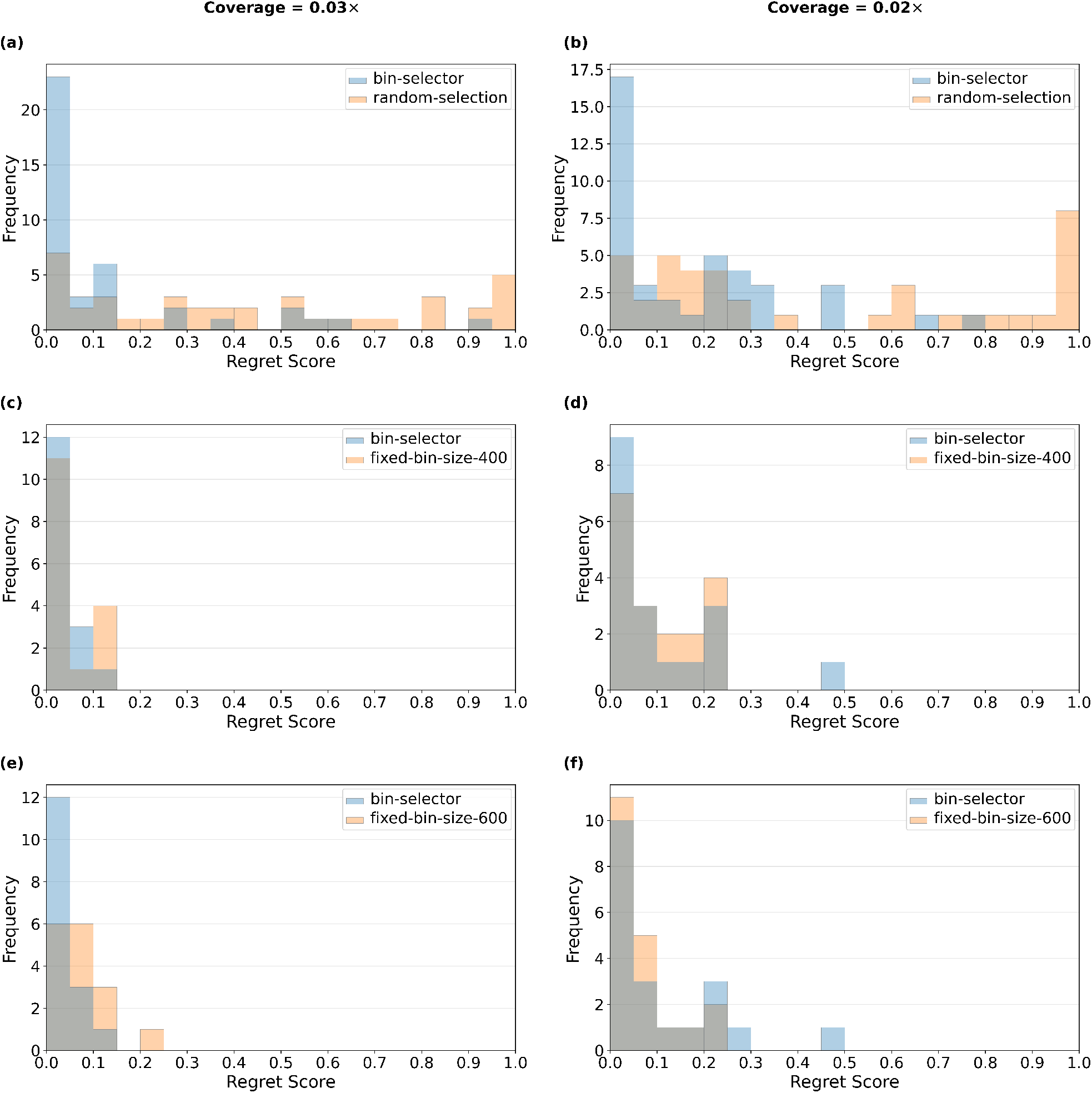
Comparison of regret scores for bin-selector and alternative approaches. Regret score comparisons of bin-selector are shown against random-selection (plots a, b), fixed-bin-size-400 (plots c, d), and fixed-bin-size-600 (plots e, f). For each row, results for sequence coverage 0.03x appear on the left and for 0.02x on the right. Each plot shows the distributions of regret scores for bin-selector and for the method being compared against, with overlapping portions of the distributions shown in gray. Smaller regret scores imply better performance. For various reasons (as detailed in the main text), the fixed-bin-size-400 and fixed-bin-size-600 approaches could not be applied to all 40 input instances for each sequence coverage level. For fair comparison, the distributions shown for bin-selector in plots c, d, e, and f are limited to the corresponding subsets of successful input instances.

When comparing against fixed-bin-size-400 and fixed-bin-size-600, we were unable to use all 40 input instances for each sequence coverage level. In particular, since SeCNV was generally unable to compute CNPs for the smaller bin sizes, including 400 and 600 kbp, we only evaluated fixed-bin-size-400 and fixed-bin-size-600 on the 20 SCOPE input instances at each coverage level. Further, since either SCOPE or MEDICC2 also sometimes failed to produce CNPs or phylogenies for 400 and/or 600 kbp bin sizes, we were able to apply the fixed-bin-size-400 approach to 16 of the 0.03x coverage input instances and 16 of the 0.02x coverage input instances, and could apply the fixed-bin-size-600 approach to 18 of the 0.03x coverage input instances and 20 of the 0.02x coverage input instances. Thus, for fairness in comparing against these approaches, we only considered the performance of bin-selector on the relevant subsets of input instances.

As the plots in the middle row of Figure 4 show, bin-selector also outperforms the fixed-bin-size-400 approach. For the 0.03x input instances, performance appears roughly matched, with bin-selector selecting a strictly better bin size than fixed-bin-size-400 in 31.25% of cases, an equally good bin size in 37.5% of cases, and a worse bin size in 31.25% of cases. However, as Figure 4(c) shows, bin-selector selects bins with smaller regret scores overall than fixed-bin-size-400, with an average regret score of 0.035 compared to 0.05 for fixed-bin-size-400. Bin-selector more clearly outperforms fixed-bin-size-400 for the 0.02x input instances, showing strictly better, equal, and worse performance in 50%, 16.67%, and 33.33% of cases, respectively.

Comparison against the fixed-bin-size-600 approach yields mixed results (bottom plots of Figure 4). In particular, we find that bin-selector substantially outperforms fixed-bin-size-600 on the 0.03x input instances, but not on the 0.02x input instances. For the 0.03x input instances bin-selector selects a strictly better bin size than fixed-bin-size-600 in 62.5% of cases, an equally good bin size in 25% of cases, and a worse bin size in 12.5% of cases. For the 0.02x coverage input instances, these values are 35%, 20%, and 45%, respectively. The strong performance of fixed-bin-size-600 on the 0.02x input instances is likely to be an artifact of the simulation parameters (such as CNA rate, mean CNA event length, etc.), used to generate the simulated datasets, which, when combined with a sequence coverage of 0.02x, result in 600 kbp being a near-ideal bin size for most of the input instances.

Overall, these results demonstrate that using the bin-selector method is likely to result in more accurate tumor cell phylogenies than simply choosing a bin size randomly or using a fixed or default bin size based on previous literature or software defaults.

### 4.3 Application of bin-selector to real datasets

We also applied bin-selector to the three real datasets. Table 3 shows the results of this application for each combination of CNP and phylogeny inference method. These results provide several interesting insights: First, we find that bin-selector generally favors larger bin sizes (1000 kbp or larger) for all three datasets. These findings are largely consistent with the trends of intra-bin NRFDs observed previously for CRC2 and T3 (Figure 1), where we found that larger bin sizes usually resulted in smaller intra-bin NRFD values, suggesting increased phylogenetic information in CNPs inferred using those bin sizes. Second, the best bin size may depend not only on the characteristics of the dataset but also on the specific inference methods used. For example, depending on the inference methods used, bin-selector suggests bin sizes ranging from 600 kbp to 3000 kbp for the CRC2 dataset. And third, despite the variability in bin sizes suggested for the same dataset based on the inference methods used, most combinations of inference methods are consistent in their support for the use of larger bin sizes for these datasets. We also note that, in the case of T5, the bin sizes selected when using SCOPE CNPs may be artificially inflated due to the previously observed, unexpectedly high similarity between the phylogenies for bin sizes 5000 and 6000 kbp (Supplementary Tables S5 andS7). Still, T5’s results with SeCNV CNPs support the use of larger bin sizes for that dataset.

**Table 3.** Results of applying bin-selector to the three real datasets. Results are shown for each of the four combinations of inference methods.

| Dataset | Inference methods | Bin size (kbp) |
| --- | --- | --- |
| CRC2 | SCOPE + DICE-star | 1500 |
| CRC2 | SCOPE + MEDICC2 | 600 |
| CRC2 | SeCNV + DICE-star | 1000 |
| CRC2 | SeCNV + MEDICC2 | 3000 |
| T3 | SCOPE + DICE-star | 600 |
| T3 | SCOPE + MEDICC2 | 3000 |
| T3 | SeCNV + DICE-star | 3000 |
| T3 | SeCNV + MEDICC2 | 3000 |
| T5 | SCOPE + DICE-star | 5000 |
| T5 | SCOPE + MEDICC2 | 5000 |
| T5 | SeCNV + DICE-star | 2500 |
| T5 | SeCNV + MEDICC2 | 2500 |

## 5 Conclusion

In this work, we demonstrated the importance of careful bin size selection when using CNPs inferred from low-coverage scDNA-seq data for tumor cell phylogeny inference, and proposed a simple computational approach for optimal bin size selection. Our analysis of the three real datasets clearly shows that even small changes in bin size can lead to drastic differences in inferred tumor cell phylogenies, and that these differences are almost always greater than can be explained by tree reconstruction error, uncertainty, or lack of phylogenetic signal in the data. We also find that these findings hold even when alternative CNP inference and tumor cell phylogeny inference methods are used. Our results using simulated datasets free from possible confounding factors further confirm these findings and clearly point to the central role of bin size in determining phylogenetic accuracy. Our evaluation of the proposed bin-selector method and other baseline approaches for bin selection shows that bin-selector is effective at identifying optimal or near-optimal bin sizes.

We view the bin-selector method as a proof-of-concept that demonstrates the feasibility of optimizing bin sizes for phylogenetic accuracy without any knowledge of the ground truth or any other evolutionary parameters. While our experiments show that bin-selector can be quite effective at selecting good bin sizes, further systematic analysis using additional simulated datasets is needed to better identify the limits of its capability. In particular, our analysis of the real datasets shows that both intra-bin and inter-bin NRFDs can be substantially higher for real datasets than what was observed for the simulated datasets used in this work, and it would be useful to bridge this gap using different and more noisy/error-prone simulated datasets. Likewise, since cancer researchers may sometimes only be interested in studying tumor evolution at the level of clones or subclones, it would be useful to conduct an in-depth assessment of how bin size selection affects clone and subclone inference and reconstruction of phylogenetic relationships between them.

It should also be possible to substantially improve upon the simple bin-selector method using more sophisticated approaches. For example, it may be possible to achieve better results by optimizing the bin size for each chromosome or large genomic region separately. Another promising approach is to use machine learning techniques to train a classifier using a variety of relevant input features, such as sequence coverage level, number of cells, intra-bin and inter-bin NRFDs, bin-selector output, etc., to estimate the best bin size.

## Data, script and code availability

An implementation of the bin-selector method is freely available open-source from GitHub at https://github.com/Tanveer14/BinSelector. All simulated and real copy number profiles, ground truth phylogenetic trees, and reconstructed phylogenetic trees used in this work are available via Zenodo at https://doi.org/10.5281/zenodo.21908781.

