## Supplementary text, figures, and tables for "Optimizing copy number profiles for tumor cell phylogeny inference from low-coverage single-cell sequencing data"

### Supplementary Material

### Supplementary Text

#### Simulation settings

To generate simulated sequencing reads, we utilized the existing software CNAsim [34] in read-generation mode (mode 2). CNAsim is designed to simulate single-cell CNPs and DNA sequencing data from tumors in a phylogenetically explicit framework. In addition to generating sequencing reads, CNAsim also reports the true phylogeny for the sampled tumor cells.

We simulated datasets with 100 cells each, of which 10% are normal cells. We varied sequencing coverage (0.02x and 0.03x) and kept most other parameter settings at their default values. Some of the parameter settings were changed from their defaults to improve overall phylogenetic resolution and focus the analysis on the effect of bin size changes. For the reference genome, we used the Genome Reference Consortium Human Build 38 (GRCh38 or Hg38), the latest available at the time. The specific CNAsim command used and list of non-default parameter settings appears below.

CNAsim command used:

```
cnasim -o <output_dir> -m 2 -r1 <reference_genome_path> -n 100 -n1 0.1 -c 5 -c2 20 -R 76 -U -v -i 2 -i1 2 -i2 3 -M -C <coverage> -B 1000000 -W 1000000 -k 10000 -l1 10000
```

Non-default parameter settings:

1. num-cells -n: 100 (total number of cells)
2. normal-fraction -n1: 0.1 (fraction of normal cells, set to 10%)
3. num-clones -c: 5 (number of clones in the population)
4. clone-mu -c2: 20 (Mean number of leaves in a subclone)
5. chrom-level-event -v: True (include chromosomal CNAs)
6. chrom-rate-super-clone -i1: 2 (Mean number of chromosomal CNAs on edges out of the founder) [Default is 1]
7. chrom-rate-clone -i2: 3 (Mean number of chromosomal CNAs on edges into clonal ancestors) [Default is 1]
8. use-uniform-coverage -M: True (Assumes uniform coverage across the genome)
9. coverage -C: 0.02 and 0.03 (Sequencing coverage across the entire genome)
10. read-length -R: 76 (derived from real datasets T3/5/8 [6])
11. use-hg38-static, U: True (Use chromosome lengths and chromosome arm ratios derived from the hg38 reference genome, excluding sex chromosomes.)
12. region-length -k: 10000 (Region length in bp. This controls the resolution at which CNA events are applied.) [Default is 1000]. We use a lower resolution for greater efficiency.

13. min-cn-length -l1: 10000 (The minimum focal CNA length in bp. This should be equal to the region-length used at minimum and no greater than the mean CNA length. We set this to the region-length used above.) [Default is 1000.]

All other parameters were kept at their default values.

### Supplementary Figures

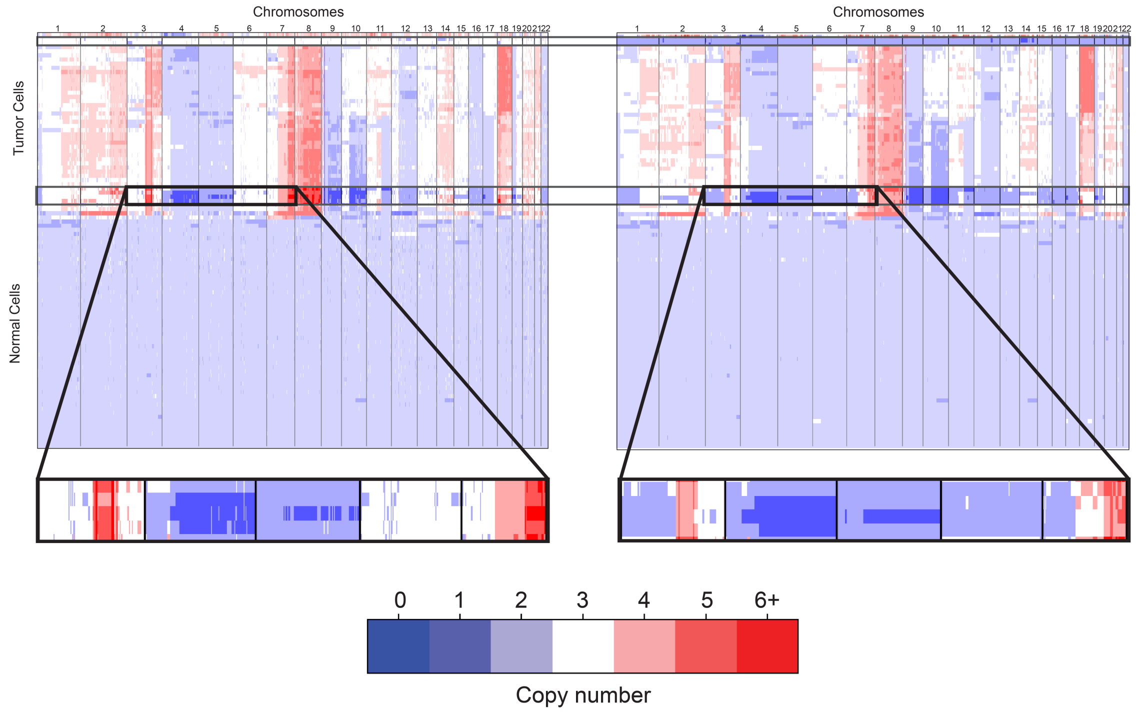


**Figure S1. CNP heatmaps of 100 cells from patient T16 [21] generated with SCOPE over two bin-sizes.** Left: 100kbp. Right: 250kbp. The tumor cell phylogenies (not shown) constructed from these two bin-sizes show different clonal compositions. As the heatmaps show, the 250kbp bins support the presence of an additional clonal population, as evidenced by unique shared mutations on chromosomes 3 and 5, whereas the 100kbp bins group those cells with an existing clonal population. Additionally, the 250kbp bins label two cells as diploid, but the 100kbp bins indicate triploid cells.

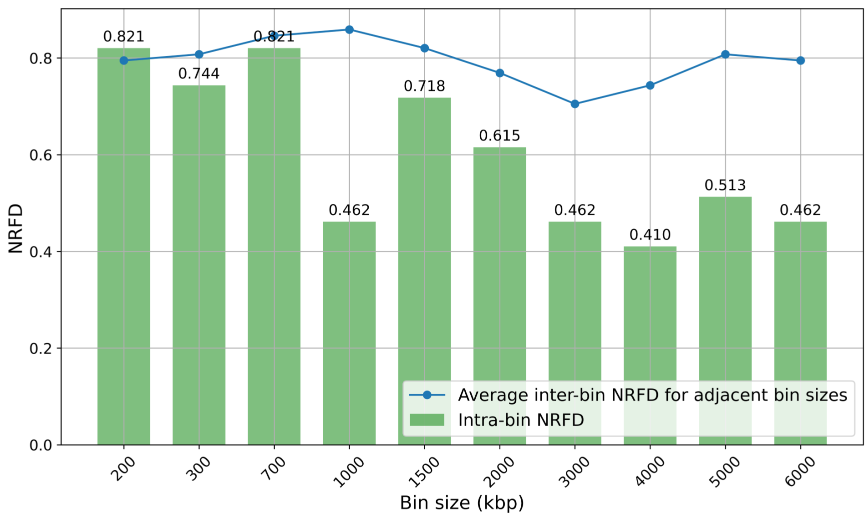


**Figure S2. Intra-bin phylogenetic distances and inter-bin distances for adjacent bins for the CRC2 dataset when using SeCNV CNPs.** All CNPs were inferred using SeCNV. For each bin size, the intra-bin distance (green bars) is the NRFD between the DICE-star phylogeny and the MEDICC2 phylogeny computed from that CNP. The inter-bin distance reported for each bin size is the average NRFD between the DICE-star phylogeny for the current bin size and the two DICE-star phylogenies for the two adjacent bin sizes; only one adjacent bin is used for the smallest and largest bin sizes.

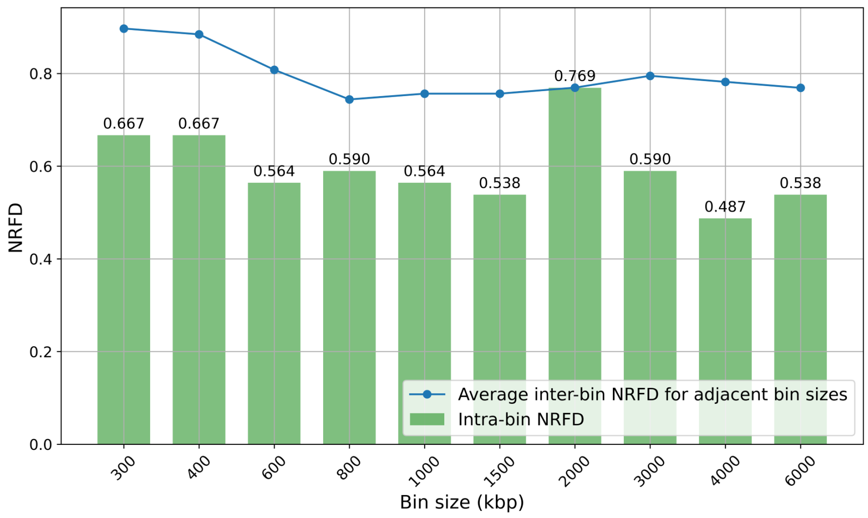


**Figure S3. Intra-bin phylogenetic distances and inter-bin distances for adjacent bins for the CRC2 dataset when using MEDICC2 phylogenies.** CNPs were inferred using SCOPE and MEDICC2 was used as the primary phylogeny reconstruction method. For each bin size, the intra-bin distance (green bars) is the NRFD between the DICE-star phylogeny and the MEDICC2 phylogeny computed from that CNP. The inter-bin distance reported for each bin size is the average NRFD between the MEDICC2 phylogeny for the current bin size and the two MEDICC2 phylogenies for the two adjacent bin sizes; only one adjacent bin is used for the smallest and largest bin sizes.

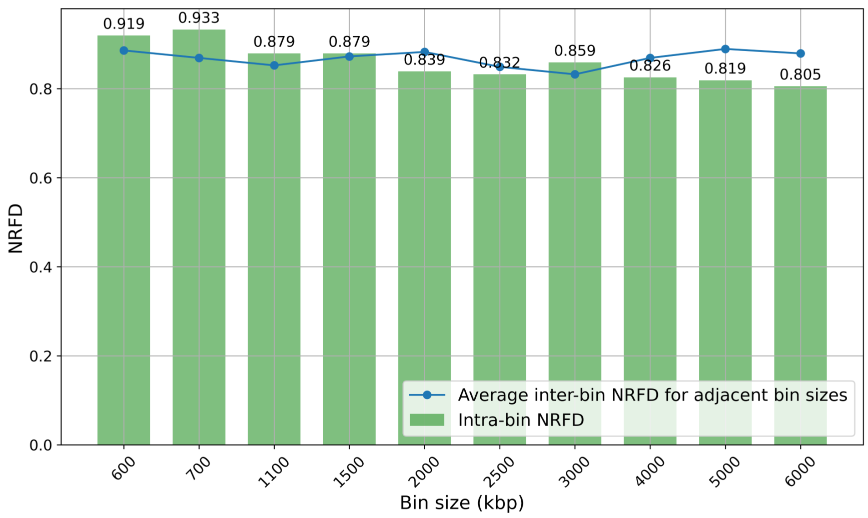


**Figure S4. Intra-bin phylogenetic distances and inter-bin distances for adjacent bins for the T3 dataset when using SeCNV CNPs.** All CNPs were inferred using SeCNV. For each bin size, the intra-bin distance (green bars) is the NRFD between the DICE-star phylogeny and the MEDICC2 phylogeny computed from that CNP. The inter-bin distance reported for each bin size is the average NRFD between the DICE-star phylogeny for the current bin size and the two DICE-star phylogenies for the two adjacent bin sizes; only one adjacent bin is used for the smallest and largest bin sizes.

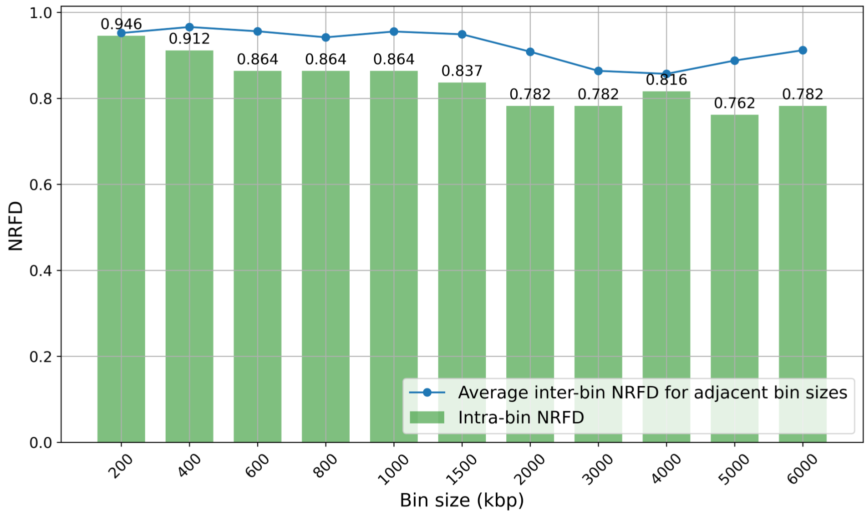


**Figure S5. Intra-bin phylogenetic distances and inter-bin distances for adjacent bins for the T3 dataset when using MEDICC2 phylogenies.** CNPs were inferred using SCOPE and MEDICC2 was used as the primary phylogeny reconstruction method. For each bin size, the intra-bin distance (green bars) is the NRFD between the DICE-star phylogeny and the MEDICC2 phylogeny computed from that CNP. The inter-bin distance reported for each bin size is the average NRFD between the MEDICC2 phylogeny for the current bin size and the two MEDICC2 phylogenies for the two adjacent bin sizes; only one adjacent bin is used for the smallest and largest bin sizes.

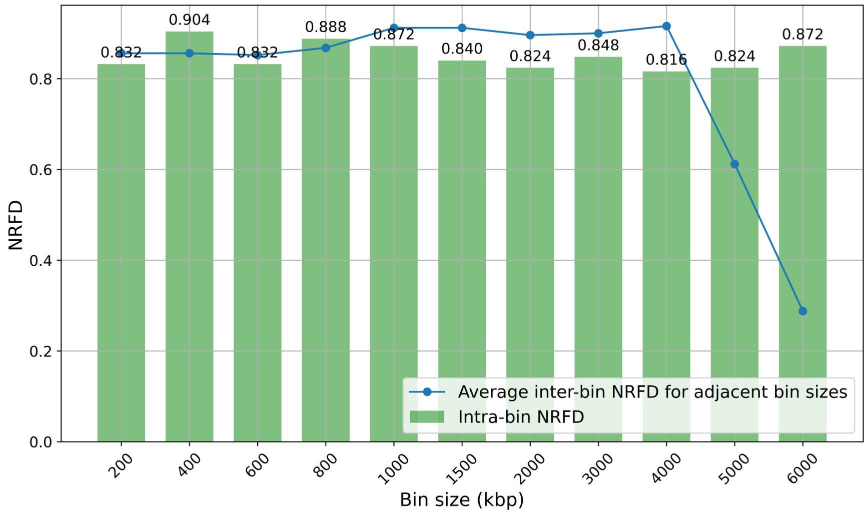


**Figure S6. Intra-bin phylogenetic distances and inter-bin distances for adjacent bins for the T5 dataset using default inference methods.** CNPs were inferred using SCOPE. For each bin size, the intra-bin distance (green bars) is the NRFD between the DICE-star phylogeny and the MEDICC2 phylogeny computed from that CNP. The inter-bin distance reported for each bin size is the average NRFD between the DICE-star phylogeny for the current bin size and the two DICE-star phylogenies for the two adjacent bin sizes; only one adjacent bin is used for the smallest and largest bin sizes.

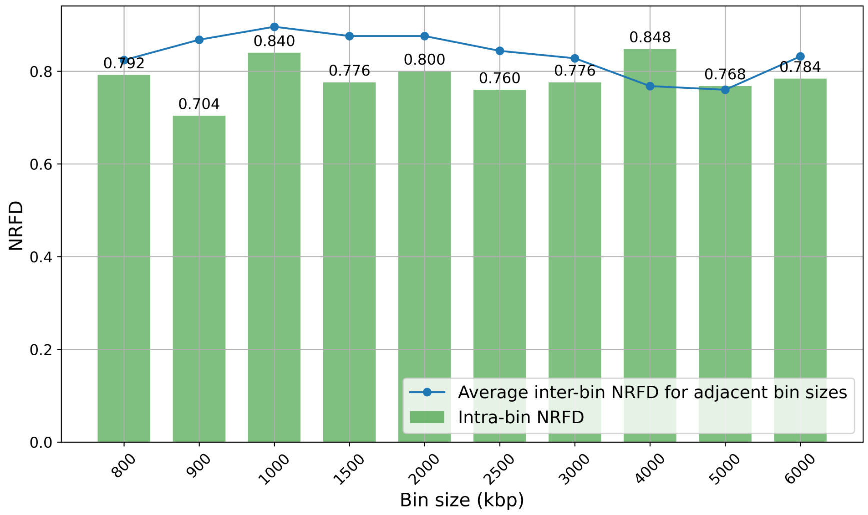


**Figure S7. Intra-bin phylogenetic distances and inter-bin distances for adjacent bins for the T5 dataset when using SeCNV CNPs.** All CNPs were inferred using SeCNV. For each bin size, the intra-bin distance (green bars) is the NRFD between the DICE-star phylogeny and the MEDICC2 phylogeny computed from that CNP. The inter-bin distance reported for each bin size is the average NRFD between the DICE-star phylogeny for the current bin size and the two DICE-star phylogenies for the two adjacent bin sizes; only one adjacent bin is used for the smallest and largest bin sizes.

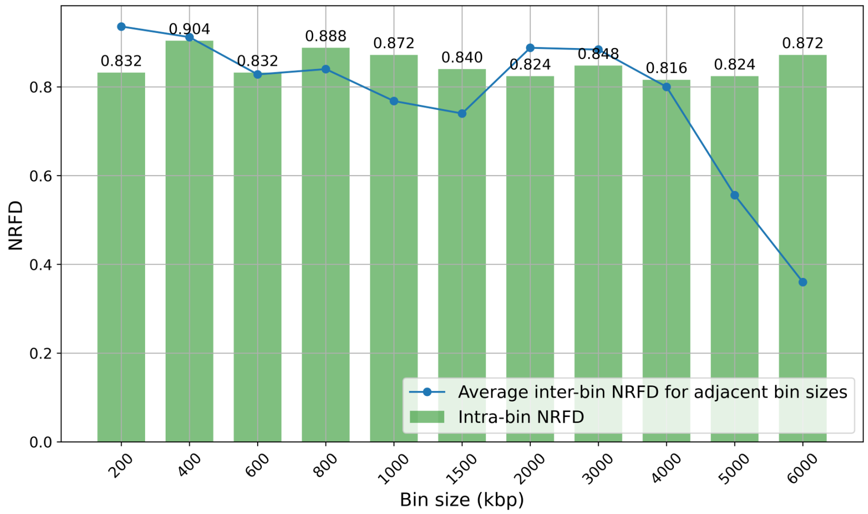


**Figure S8. Intra-bin phylogenetic distances and inter-bin distances for adjacent bins for the T5 dataset when using MEDICC2 phylogenies.** CNPs were inferred using SCOPE and MEDICC2 was used as the primary phylogeny reconstruction method. For each bin size, the intra-bin distance (green bars) is the NRFD between the DICE-star phylogeny and the MEDICC2 phylogeny computed from that CNP. The inter-bin distance reported for each bin size is the average NRFD between the MEDICC2 phylogeny for the current bin size and the two MEDICC2 phylogenies for the two adjacent bin sizes; only one adjacent bin is used for the smallest and largest bin sizes.

#### Supplementary Tables

**Table S1. Inter-bin distance matrix for tumor cell phylogenies inferred from CNPs of different bin sizes for the CRC2 dataset using SeCNV and DICE-star.** The distance matrix is symmetric, phylogenetic distances are reported in NRFD, and all bin sizes are in kbp. CNPs were inferred using SeCNV and phylogenies were reconstructed using DICE-star. NRFDs between adjacent bin sizes appear shaded in blue. Note that a bin size of 300 kbp was used instead of 400 kbp, and 700 kbp was used instead of bin sizes 600 and 800 kbp, since SeCNV failed to compute CNPs for this dataset for 400, 600, and 800 kbp.

| Bin size | 200 | 300 | 700 | 1000 | 1500 | 2000 | 3000 | 4000 | 5000 | 6000 |
|---|---|---|---|---|---|---|---|---|---|---|
| 200 | 0.000 | 0.795 | 0.846 | 0.821 | 0.897 | 0.821 | 0.846 | 0.923 | 0.923 | 0.846 |
| 300 | 0.795 | 0.000 | 0.821 | 0.821 | 0.949 | 0.846 | 0.872 | 0.897 | 0.846 | 0.795 |
| 700 | 0.846 | 0.821 | 0.000 | 0.872 | 0.923 | 0.846 | 0.923 | 0.949 | 0.949 | 0.872 |
| 1000 | 0.821 | 0.821 | 0.872 | 0.000 | 0.846 | 0.667 | 0.718 | 0.744 | 0.872 | 0.769 |
| 1500 | 0.897 | 0.949 | 0.923 | 0.846 | 0.000 | 0.795 | 0.769 | 0.795 | 0.821 | 0.923 |
| 2000 | 0.821 | 0.846 | 0.846 | 0.667 | 0.795 | 0.000 | 0.744 | 0.744 | 0.872 | 0.846 |
| 3000 | 0.846 | 0.872 | 0.923 | 0.718 | 0.769 | 0.744 | 0.000 | 0.667 | 0.821 | 0.846 |
| 4000 | 0.923 | 0.897 | 0.949 | 0.744 | 0.795 | 0.744 | 0.667 | 0.000 | 0.821 | 0.897 |
| 5000 | 0.923 | 0.846 | 0.949 | 0.872 | 0.821 | 0.872 | 0.821 | 0.821 | 0.000 | 0.795 |
| 6000 | 0.846 | 0.795 | 0.872 | 0.769 | 0.923 | 0.846 | 0.846 | 0.897 | 0.795 | 0.000 |

**Table S2. Inter-bin distance matrix for tumor cell phylogenies inferred from CNPs of different bin sizes for the CRC2 dataset using SCOPE and MEDICC2.** The distance matrix is symmetric, phylogenetic distances are reported in NRFD, and all bin sizes are in kbp. CNPs were inferred using SCOPE and phylogenies were reconstructed using MEDICC2. NRFDs between adjacent bin sizes appear shaded in blue. Note that a bin size of 300 kbp was used instead of 200 kbp and that results are not shown for bin size 5000 kbp since SCOPE failed to compute CNPs for this dataset for 200 kbp and 5000 kbp.

| Bin size | 300 | 400 | 600 | 800 | 1000 | 1500 | 2000 | 3000 | 4000 | 6000 |
|---|---|---|---|---|---|---|---|---|---|---|
| 300 | 0.000 | 0.897 | 0.821 | 0.923 | 0.872 | 0.897 | 0.923 | 0.949 | 0.872 | 0.897 |
| 400 | 0.897 | 0.000 | 0.872 | 0.872 | 0.872 | 0.923 | 0.872 | 0.949 | 0.897 | 0.897 |
| 600 | 0.821 | 0.872 | 0.000 | 0.744 | 0.769 | 0.667 | 0.795 | 0.872 | 0.744 | 0.821 |
| 800 | 0.923 | 0.872 | 0.744 | 0.000 | 0.744 | 0.795 | 0.846 | 0.846 | 0.821 | 0.846 |
| 1000 | 0.872 | 0.872 | 0.769 | 0.744 | 0.000 | 0.769 | 0.872 | 0.897 | 0.795 | 0.872 |
| 1500 | 0.897 | 0.923 | 0.667 | 0.795 | 0.769 | 0.000 | 0.744 | 0.744 | 0.821 | 0.872 |
| 2000 | 0.923 | 0.872 | 0.795 | 0.846 | 0.872 | 0.744 | 0.000 | 0.795 | 0.667 | 0.795 |
| 3000 | 0.949 | 0.949 | 0.872 | 0.846 | 0.897 | 0.744 | 0.795 | 0.000 | 0.795 | 0.846 |
| 4000 | 0.872 | 0.897 | 0.744 | 0.821 | 0.795 | 0.821 | 0.667 | 0.795 | 0.000 | 0.769 |
| 6000 | 0.897 | 0.897 | 0.821 | 0.846 | 0.872 | 0.872 | 0.795 | 0.846 | 0.769 | 0.000 |

**Table S3. Inter-bin distance matrix for tumor cell phylogenies inferred from CNPs of different bin sizes for the T3 dataset using SeCNV and DICE-star.** The distance matrix is symmetric, phylogenetic distances are reported in NRFD, and all bin sizes are in kbp. CNPs were inferred using SeCNV and phylogenies were reconstructed using DICE-star. NRFDs between adjacent bin sizes appear shaded in blue. Note that bin sizes 700 and 1100 kbp were used instead of 800 and 1000 kbp, respectively. SeCNV failed to run for bin sizes smaller than 600 kbp and on bin sizes 800 and 1000 kbp on this dataset. An additional bin size, 2500 kbp, was used to make up for the large number of missing bin sizes.

| Bin size | 600 | 700 | 1100 | 1500 | 2000 | 2500 | 3000 | 4000 | 5000 | 6000 |
|---|---|---|---|---|---|---|---|---|---|---|
| 600 | 0.000 | 0.886 | 0.852 | 0.873 | 0.859 | 0.899 | 0.886 | 0.893 | 0.913 | 0.926 |
| 700 | 0.886 | 0.000 | 0.852 | 0.873 | 0.893 | 0.866 | 0.879 | 0.946 | 0.906 | 0.920 |
| 1100 | 0.852 | 0.852 | 0.000 | 0.852 | 0.859 | 0.866 | 0.846 | 0.920 | 0.913 | 0.899 |
| 1500 | 0.873 | 0.873 | 0.852 | 0.000 | 0.893 | 0.893 | 0.873 | 0.906 | 0.913 | 0.886 |
| 2000 | 0.859 | 0.893 | 0.859 | 0.893 | 0.000 | 0.873 | 0.899 | 0.906 | 0.920 | 0.866 |
| 2500 | 0.899 | 0.866 | 0.866 | 0.893 | 0.873 | 0.000 | 0.826 | 0.906 | 0.920 | 0.886 |
| 3000 | 0.886 | 0.879 | 0.846 | 0.873 | 0.899 | 0.826 | 0.000 | 0.839 | 0.873 | 0.866 |
| 4000 | 0.893 | 0.946 | 0.920 | 0.906 | 0.906 | 0.906 | 0.839 | 0.000 | 0.899 | 0.913 |
| 5000 | 0.913 | 0.906 | 0.913 | 0.913 | 0.920 | 0.920 | 0.873 | 0.899 | 0.000 | 0.879 |
| 6000 | 0.926 | 0.920 | 0.899 | 0.886 | 0.866 | 0.886 | 0.866 | 0.913 | 0.879 | 0.000 |

**Table S4. Inter-bin distance matrix for tumor cell phylogenies inferred from CNPs of different bin sizes for the T3 dataset using SCOPE and MEDICC2.** The distance matrix is symmetric, phylogenetic distances are reported in NRFD, and all bin sizes are in kbp. CNPs were inferred using SCOPE and phylogenies were reconstructed using MEDICC2. NRFDs between adjacent bin sizes appear shaded in blue.

| Bin size | 200 | 400 | 600 | 800 | 1000 | 1500 | 2000 | 3000 | 4000 | 5000 | 6000 |
|---|---|---|---|---|---|---|---|---|---|---|---|
| 200 | 0.000 | 0.952 | 0.959 | 0.946 | 0.986 | 0.986 | 0.973 | 0.966 | 0.946 | 0.973 | 0.973 |
| 400 | 0.952 | 0.000 | 0.980 | 0.980 | 0.959 | 0.980 | 0.952 | 0.959 | 0.952 | 0.986 | 0.966 |
| 600 | 0.959 | 0.980 | 0.000 | 0.932 | 0.986 | 0.966 | 0.980 | 0.973 | 0.973 | 0.959 | 0.966 |
| 800 | 0.946 | 0.980 | 0.932 | 0.000 | 0.952 | 0.973 | 0.966 | 0.959 | 0.973 | 0.980 | 0.973 |
| 1000 | 0.986 | 0.959 | 0.986 | 0.952 | 0.000 | 0.959 | 0.966 | 0.966 | 0.980 | 0.973 | 0.980 |
| 1500 | 0.986 | 0.980 | 0.966 | 0.973 | 0.959 | 0.000 | 0.939 | 0.952 | 0.939 | 0.959 | 0.980 |
| 2000 | 0.973 | 0.952 | 0.980 | 0.966 | 0.966 | 0.939 | 0.000 | 0.878 | 0.912 | 0.946 | 0.932 |
| 3000 | 0.966 | 0.959 | 0.973 | 0.959 | 0.966 | 0.952 | 0.878 | 0.000 | 0.850 | 0.857 | 0.871 |
| 4000 | 0.946 | 0.952 | 0.973 | 0.973 | 0.980 | 0.939 | 0.912 | 0.850 | 0.000 | 0.864 | 0.946 |
| 5000 | 0.973 | 0.986 | 0.959 | 0.980 | 0.973 | 0.959 | 0.946 | 0.857 | 0.864 | 0.000 | 0.912 |
| 6000 | 0.973 | 0.966 | 0.966 | 0.973 | 0.980 | 0.980 | 0.932 | 0.871 | 0.946 | 0.912 | 0.000 |

**Table S5. Inter-bin distance matrix for tumor cell phylogenies inferred from CNPs of different bin sizes for the T5 dataset using default inference methods.** The distance matrix is symmetric, phylogenetic distances are reported in NRFD, and all bin sizes are in kbp. CNPs were inferred using SCOPE and phylogenies were reconstructed using DICE-star. NRFDs between adjacent bin sizes appear shaded in blue.

| Bin size | 200 | 400 | 600 | 800 | 1000 | 1500 | 2000 | 3000 | 4000 | 5000 | 6000 |
|---|---|---|---|---|---|---|---|---|---|---|---|
| 200 | 0.000 | 0.856 | 0.856 | 0.848 | 0.880 | 0.928 | 0.896 | 0.952 | 0.968 | 0.960 | 0.984 |
| 400 | 0.856 | 0.000 | 0.856 | 0.896 | 0.872 | 0.920 | 0.896 | 0.928 | 0.944 | 0.960 | 0.984 |
| 600 | 0.856 | 0.856 | 0.000 | 0.848 | 0.896 | 0.912 | 0.904 | 0.944 | 0.952 | 0.960 | 0.968 |
| 800 | 0.848 | 0.896 | 0.848 | 0.000 | 0.888 | 0.920 | 0.896 | 0.944 | 0.952 | 0.960 | 0.968 |
| 1000 | 0.880 | 0.872 | 0.896 | 0.888 | 0.000 | 0.936 | 0.896 | 0.920 | 0.936 | 0.968 | 0.992 |
| 1500 | 0.928 | 0.920 | 0.912 | 0.920 | 0.936 | 0.000 | 0.888 | 0.912 | 0.944 | 0.976 | 0.984 |
| 2000 | 0.896 | 0.896 | 0.904 | 0.896 | 0.896 | 0.888 | 0.000 | 0.904 | 0.912 | 0.928 | 0.960 |
| 3000 | 0.952 | 0.928 | 0.944 | 0.944 | 0.920 | 0.912 | 0.904 | 0.000 | 0.896 | 0.936 | 0.960 |
| 4000 | 0.968 | 0.944 | 0.952 | 0.952 | 0.936 | 0.944 | 0.912 | 0.896 | 0.000 | 0.936 | 0.944 |
| 5000 | 0.960 | 0.960 | 0.960 | 0.960 | 0.968 | 0.976 | 0.928 | 0.936 | 0.936 | 0.000 | 0.288 |
| 6000 | 0.984 | 0.984 | 0.968 | 0.968 | 0.992 | 0.984 | 0.960 | 0.960 | 0.944 | 0.288 | 0.000 |

**Table S6. Inter-bin distance matrix for tumor cell phylogenies inferred from CNPs of different bin sizes for the T5 dataset using SeCNV and DICE-star.** The distance matrix is symmetric, phylogenetic distances are reported in NRFD, and all bin sizes are in kbp. CNPs were inferred using SeCNV and phylogenies were reconstructed using DICE-star. NRFDs between adjacent bin sizes appear shaded in blue. SeCNV failed to run for bin sizes smaller than 800 kbp. Additional bin sizes 900 and 2500 kbp were used to make up for the large number of missing bin sizes.

| Bin size | 800 | 900 | 1000 | 1500 | 2000 | 2500 | 3000 | 4000 | 5000 | 6000 |
|---|---|---|---|---|---|---|---|---|---|---|
| 800 | 0.000 | 0.824 | 0.904 | 0.888 | 0.896 | 0.872 | 0.872 | 0.904 | 0.896 | 0.912 |
| 900 | 0.824 | 0.000 | 0.912 | 0.912 | 0.904 | 0.888 | 0.904 | 0.896 | 0.904 | 0.920 |
| 1000 | 0.904 | 0.912 | 0.000 | 0.880 | 0.896 | 0.904 | 0.912 | 0.920 | 0.920 | 0.936 |
| 1500 | 0.888 | 0.912 | 0.880 | 0.000 | 0.872 | 0.816 | 0.824 | 0.856 | 0.872 | 0.872 |
| 2000 | 0.896 | 0.904 | 0.896 | 0.872 | 0.000 | 0.880 | 0.888 | 0.904 | 0.896 | 0.880 |
| 2500 | 0.872 | 0.888 | 0.904 | 0.816 | 0.880 | 0.000 | 0.808 | 0.608 | 0.776 | 0.856 |
| 3000 | 0.872 | 0.904 | 0.912 | 0.824 | 0.888 | 0.808 | 0.000 | 0.848 | 0.864 | 0.824 |
| 4000 | 0.904 | 0.896 | 0.920 | 0.856 | 0.904 | 0.608 | 0.848 | 0.000 | 0.688 | 0.832 |
| 5000 | 0.896 | 0.904 | 0.920 | 0.872 | 0.896 | 0.776 | 0.864 | 0.688 | 0.000 | 0.832 |
| 6000 | 0.912 | 0.920 | 0.936 | 0.872 | 0.880 | 0.856 | 0.824 | 0.832 | 0.832 | 0.000 |

**Table S7. Inter-bin distance matrix for tumor cell phylogenies inferred from CNPs of different bin sizes for the T5 dataset using SCOPE and MEDICC2.** The distance matrix is symmetric, phylogenetic distances are reported in NRFD, and all bin sizes are in kbp. CNPs were inferred using SCOPE and phylogenies were reconstructed using MEDICC2. NRFDs between adjacent bin sizes appear shaded in blue.

| Bin size | 200 | 400 | 600 | 800 | 1000 | 1500 | 2000 | 3000 | 4000 | 5000 | 6000 |
|---|---|---|---|---|---|---|---|---|---|---|---|
| 200 | 0.000 | 0.936 | 0.928 | 0.952 | 0.960 | 0.968 | 0.952 | 0.976 | 0.952 | 0.952 | 0.960 |
| 400 | 0.936 | 0.000 | 0.888 | 0.888 | 0.864 | 0.904 | 0.896 | 0.912 | 0.912 | 0.928 | 0.928 |
| 600 | 0.928 | 0.888 | 0.000 | 0.768 | 0.872 | 0.856 | 0.848 | 0.872 | 0.808 | 0.880 | 0.880 |
| 800 | 0.952 | 0.888 | 0.768 | 0.000 | 0.912 | 0.920 | 0.776 | 0.904 | 0.840 | 0.792 | 0.792 |
| 1000 | 0.960 | 0.864 | 0.872 | 0.912 | 0.000 | 0.624 | 0.856 | 0.736 | 0.776 | 0.864 | 0.872 |
| 1500 | 0.968 | 0.904 | 0.856 | 0.920 | 0.624 | 0.000 | 0.856 | 0.680 | 0.768 | 0.880 | 0.880 |
| 2000 | 0.952 | 0.896 | 0.848 | 0.776 | 0.856 | 0.856 | 0.000 | 0.920 | 0.792 | 0.632 | 0.656 |
| 3000 | 0.976 | 0.912 | 0.872 | 0.904 | 0.736 | 0.680 | 0.920 | 0.000 | 0.848 | 0.888 | 0.888 |
| 4000 | 0.952 | 0.912 | 0.808 | 0.840 | 0.776 | 0.768 | 0.792 | 0.848 | 0.000 | 0.752 | 0.712 |
| 5000 | 0.952 | 0.928 | 0.880 | 0.792 | 0.864 | 0.880 | 0.632 | 0.888 | 0.752 | 0.000 | 0.360 |
| 6000 | 0.960 | 0.928 | 0.880 | 0.792 | 0.872 | 0.880 | 0.656 | 0.888 | 0.712 | 0.360 | 0.000 |

**Table S8. Inter-bin distance matrix for tumor cell phylogenies inferred from CNPs of different bin sizes for the 0.03x sequence coverage simulated datasets when SeCNV is used for CNP inference.** The distance matrix is symmetric, phylogenetic distances are reported in NRFD, and all bin sizes are in kbp. Datasets were simulated with 0.03x coverage. CNPs were inferred using SeCNV and phylogenies were reconstructed using DICE-star. NRFDs between adjacent bin sizes appear shaded in blue. All NRFDs are averaged over 10 simulated datasets. Some inter-bin NRFDs are missing since SeCNV could not compute CNPs for at least one of the bin sizes involved in those inter-bin calculations for any of the 10 replicates.

| | **200** | **400** | **600** | **800** | **1000** | **1500** | **2000** | **3000** | **4000** | **5000** | **6000** |
|---|---|---|---|---|---|---|---|---|---|---|---|
| 200 | 0.000 | – | 0.619 | – | 0.650 | 0.670 | 0.691 | 0.732 | 0.784 | 0.773 | 0.835 |
| 400 | – | 0.000 | 0.567 | 0.557 | 0.577 | 0.546 | 0.598 | 0.660 | 0.660 | 0.691 | 0.701 |
| 600 | 0.619 | 0.567 | 0.000 | 0.445 | 0.486 | 0.495 | 0.552 | 0.615 | 0.658 | 0.684 | 0.716 |
| 800 | – | 0.557 | 0.445 | 0.000 | 0.527 | 0.524 | 0.551 | 0.608 | 0.649 | 0.685 | 0.716 |
| 1000 | 0.650 | 0.577 | 0.486 | 0.527 | 0.000 | 0.427 | 0.484 | 0.597 | 0.618 | 0.664 | 0.709 |
| 1500 | 0.670 | 0.546 | 0.495 | 0.524 | 0.427 | 0.000 | 0.450 | 0.574 | 0.608 | 0.659 | 0.694 |
| 2000 | 0.691 | 0.598 | 0.552 | 0.551 | 0.484 | 0.450 | 0.000 | 0.560 | 0.597 | 0.649 | 0.689 |
| 3000 | 0.732 | 0.660 | 0.615 | 0.608 | 0.597 | 0.574 | 0.560 | 0.000 | 0.549 | 0.586 | 0.650 |
| 4000 | 0.784 | 0.660 | 0.658 | 0.649 | 0.618 | 0.608 | 0.597 | 0.549 | 0.000 | 0.592 | 0.645 |
| 5000 | 0.773 | 0.691 | 0.684 | 0.685 | 0.664 | 0.659 | 0.649 | 0.586 | 0.592 | 0.000 | 0.648 |
| 6000 | 0.835 | 0.701 | 0.716 | 0.716 | 0.709 | 0.694 | 0.689 | 0.650 | 0.645 | 0.648 | 0.000 |

**Table S9. Inter-bin distance matrix for tumor cell phylogenies inferred from CNPs of different bin sizes for the 0.03x sequence coverage simulated datasets using MEDICC2 for phylogeny reconstruction.** The distance matrix is symmetric, phylogenetic distances are reported in NRFD, and all bin sizes are in kbp. Datasets were simulated with 0.03x coverage. CNPs were inferred using SCOPE and phylogenies were reconstructed using MEDICC2. NRFDs between adjacent bin sizes appear shaded in blue. All NRFDs are averaged over 10 simulated datasets.

| Bin size | **200** | **400** | **600** | **800** | **1000** | **1500** | **2000** | **3000** | **4000** | **5000** | **6000** |
|---|---|---|---|---|---|---|---|---|---|---|---|
| 200 | 0.000 | 0.486 | 0.492 | 0.501 | 0.524 | 0.568 | 0.607 | 0.692 | 0.732 | 0.739 | 0.741 |
| 400 | 0.486 | 0.000 | 0.307 | 0.317 | 0.334 | 0.410 | 0.500 | 0.619 | 0.687 | 0.704 | 0.694 |
| 600 | 0.492 | 0.307 | 0.000 | 0.255 | 0.307 | 0.379 | 0.462 | 0.613 | 0.682 | 0.706 | 0.696 |
| 800 | 0.501 | 0.317 | 0.255 | 0.000 | 0.237 | 0.343 | 0.431 | 0.595 | 0.675 | 0.692 | 0.688 |
| 1000 | 0.524 | 0.334 | 0.307 | 0.237 | 0.000 | 0.326 | 0.438 | 0.602 | 0.679 | 0.692 | 0.698 |
| 1500 | 0.568 | 0.410 | 0.379 | 0.343 | 0.326 | 0.000 | 0.385 | 0.561 | 0.658 | 0.692 | 0.684 |
| 2000 | 0.607 | 0.500 | 0.462 | 0.431 | 0.438 | 0.385 | 0.000 | 0.576 | 0.648 | 0.683 | 0.685 |
| 3000 | 0.692 | 0.619 | 0.613 | 0.595 | 0.602 | 0.561 | 0.576 | 0.000 | 0.633 | 0.673 | 0.700 |
| 4000 | 0.732 | 0.687 | 0.682 | 0.675 | 0.679 | 0.658 | 0.648 | 0.633 | 0.000 | 0.618 | 0.647 |
| 5000 | 0.739 | 0.704 | 0.706 | 0.692 | 0.692 | 0.692 | 0.683 | 0.673 | 0.618 | 0.000 | 0.544 |
| 6000 | 0.741 | 0.694 | 0.696 | 0.688 | 0.698 | 0.684 | 0.685 | 0.700 | 0.647 | 0.544 | 0.000 |
